# CFM: Confinement Force Microscopy—a dynamic, precise and stable microconfiner for traction force microscopy in spatial confinement

**DOI:** 10.1101/2023.08.22.554088

**Authors:** Fatemeh Abbasi, Katharina Rieck, Matthias Brandt, Maja Matis, Eva Kiermaier, Timo Betz

## Abstract

Cells migrating through tissues experience changing physical confinement, yet methods to dynamically control confinement while quantifying the resulting forces remain limited. Here, we present a microconfiner platform for live-cell imaging that enables programmable confinement, allowing real-time control over the level, timing and frequency of confinement while measuring traction forces exerted on the microenvironment, a method we term confinement force microscopy (CFM). Using CFM, we find that cells respond to confinement in two phases: a rapid passive stress rise caused by compression of the cell body and nucleus against the substrate, followed by an active stress increase associated with enhanced contractility, intracellular pressure buildup and bleb formation. Bleb expansion can partially relieve pressure and reduce stress on the surroundings. ROCK and myosin II inhibition both reduce stress generation, but with distinct effects on blebbing. Overall, CFM provides a versatile approach to study dynamic mechanical adaptation in tissue-like environments.

## 1 Introduction

Cells in living tissues experience complex and dynamic physical environments. Unlike the simplified and often planar conditions used *in vitro*, cells *in vivo* are subject to mechanical constraints and geometric confinement that influence many aspects of their behavior^1–4^. During development, immune surveillance, and tumor invasion, cells encounter boundaries imposed by the extracellular matrix (ECM), neighboring cells, or tissue architecture^5–8^. These interactions affect signaling, cytoskeletal organization, nuclear morphology, migration, and fate decisions^9–13^.

Spatial confinement is a key biomechanical input because it acts not only as a passive boundary but also as an active regulator of cellular mechanics. Immune cells repeatedly deform while traversing constricted tissue spaces^14^, and epithelial or cancer cells experience confinement from neighboring cells during collective migration^2,15^. Confinement can alter intracellular pressure^16^, nuclear shape^17–19^, and force generation^20^, with important consequences for cell function.

A wide range of platforms has been developed to study cellular responses to confinement. These include microchannels with defined dimensions and topography^11,21–25^, collagen or fibrin matrices with tunable density^9,26^, parallel plate compression devices with fixed spacings^10,20,27,28^, as well as commercial instruments such as atomic force microscopy (AFM)^17,29^, CellScale MicroSquisher^30,31^ and related cytocompression setups^32,33^. These systems have provided important insights into nuclear mechanics, cytoskeletal rearrangements, and migratory strategies^9–11,17,18,26,27,34–36^, but they often involve trade-offs between imaging access, dynamic control, and force quantification (see Supplementary Table 1).

Here, we introduce Confinement Force Microscopy (CFM), a platform that enables dynamic vertical confinement of live cells combined with three-dimensional, post hoc quantification of both lateral and normal cellular stresses. Compared with many previous approaches, CFM permits time-resolved confinement while preserving imaging access and allows downstream analysis of cellular force responses from the acquired stress maps.

Using CFM, we observed a biphasic response to compression: an initial rapid stress spike consistent with passive mechanics was followed by a slower, actomyosin-dependent increase in stress accompanied by adaptive bleb formation. Contractility inhibition reduced overall stress generation, but its effects on blebbing differed between perturbations: ROCK inhibition completely abolished blebbing, whereas blebbistatin-treated cells could still bleb under high confinement, despite lacking the sustained stress increase observed in control cells.

Furthermore, we subjected single HeLa cells to periodically modulated confinement with controlled amplitude and timing. The resulting stress profiles demonstrate the potential to track changes in single-cell viscoelasticity under repeated loading and to study fatigue-like adaptation and history-dependent force generation.

Together, these findings establish CFM as a versatile tool for probing how cells sense, resist and adapt to mechanical compression.

## 2 Results

### 2.1 Design of CFM

CFM is composed of a piezo-driven, motorized linear stage (U-521 PILine^®^ Linear Stage with a C-867.1U PILine^®^ Motion Controller; PI, Germany) and a capacitive displacement sensor (CSH1-CAm1,4 with a capaNCDT 6110 single-channel capacitive measuring system; MICRO-EPSILON, Germany) (Fig. 1a–d). The piezo stage allows dynamic adjustment of the confinement, from millimeters down to 0.5 *µ*m, thus covering the working distance of common objectives. As both cell size and spreading vary not only across cell types but also during experiments, the degree of confinement relative to cellular dimensions is cell-dependent even when the distance between the substrates is fixed. Here, the motorized stage allows easy, real-time adjustment of the confinement level during live imaging, based on the experimental conditions, and enables imaging and recording of cellular behaviors such as migration and changes in cell area at the desired confinement level. Additionally, the sensor provides long-term stability of the confinement level (experimentally tested for up to 20 h) via closed-loop feedback with the piezo motor (Fig. 1e-f). After adjusting the distance between the substrates, even a 500 nm change in sensor position will trigger motor movement to compensate. The CFM instrument consists of a metal body (EN 1.4122, X39CrMo17-1 martensitic stainless steel for the sensor holder and 6061-T6 aluminium alloy for the remaining structural components) (Fig. 1a, number 1), which is designed to be inserted in standard microscope stages for imaging (Fig. 1d). The upper assembly of the confiner consists of a lever arm (Fig. 1a, number 2) and a cylinder (Fig. 1a-b, number 3) to which the top coverslip carrying the top polyacrylamide (PAA) hydrogel (Fig. 1c, number 4) is glued. The position of this arm is adjusted by the piezo motor (Fig. 1a, number 5), which moves the top gel up and down. A custom LabVIEW program available on Zenodo (DOI: 10.5281/zenodo.19535276) continuously reads the signal from the capacitive sensor mounted on the lever arm (Fig. 1a, number 6) and uses this feedback to control piezo movement in a closed loop, thereby setting the top gel to the desired position. A small metal cylinder positioned beneath the sensor serves as the conductive target for capacitive distance measurement (Fig. 1a, number 7), helping to maintain a stable distance between the top gel (Fig. 1c, number 8) and the bottom gel (Fig. 1c, number 9). The bottom PAA gel is polymerized on a glass-bottom Petri dish (Fig. 1b, number 10), which is placed in the open area of the CFM bottom plate (Fig. 1a, red circle). The Petri dish is secured with two clamps (Fig. 1a, number 11), which are screwed on the CFM body. The mechanical stability of the glue is crucial for the stability of the CFM and can be achieved by a UV-activated glue (Norland Optical Adhesive 107, Thorlabs, Germany) that can be washed away with warm water after the experiment to reuse the metal cylinder. The cylinder is made of two materials: acrylic (Plexiglas) for compatibility with bright-field microscopy and stainless steel for more stable fluorescence microscopy. Cells are seeded either on the bottom or the top gel, depending on the experiment (Fig. 1c). The bottom plane of the CFM (Fig. 1a, number 1) has the same dimensions as a standard 6-well dish, which allows insertion into microscope stages for live-cell imaging (Fig. 1d).

**Figure 1.**
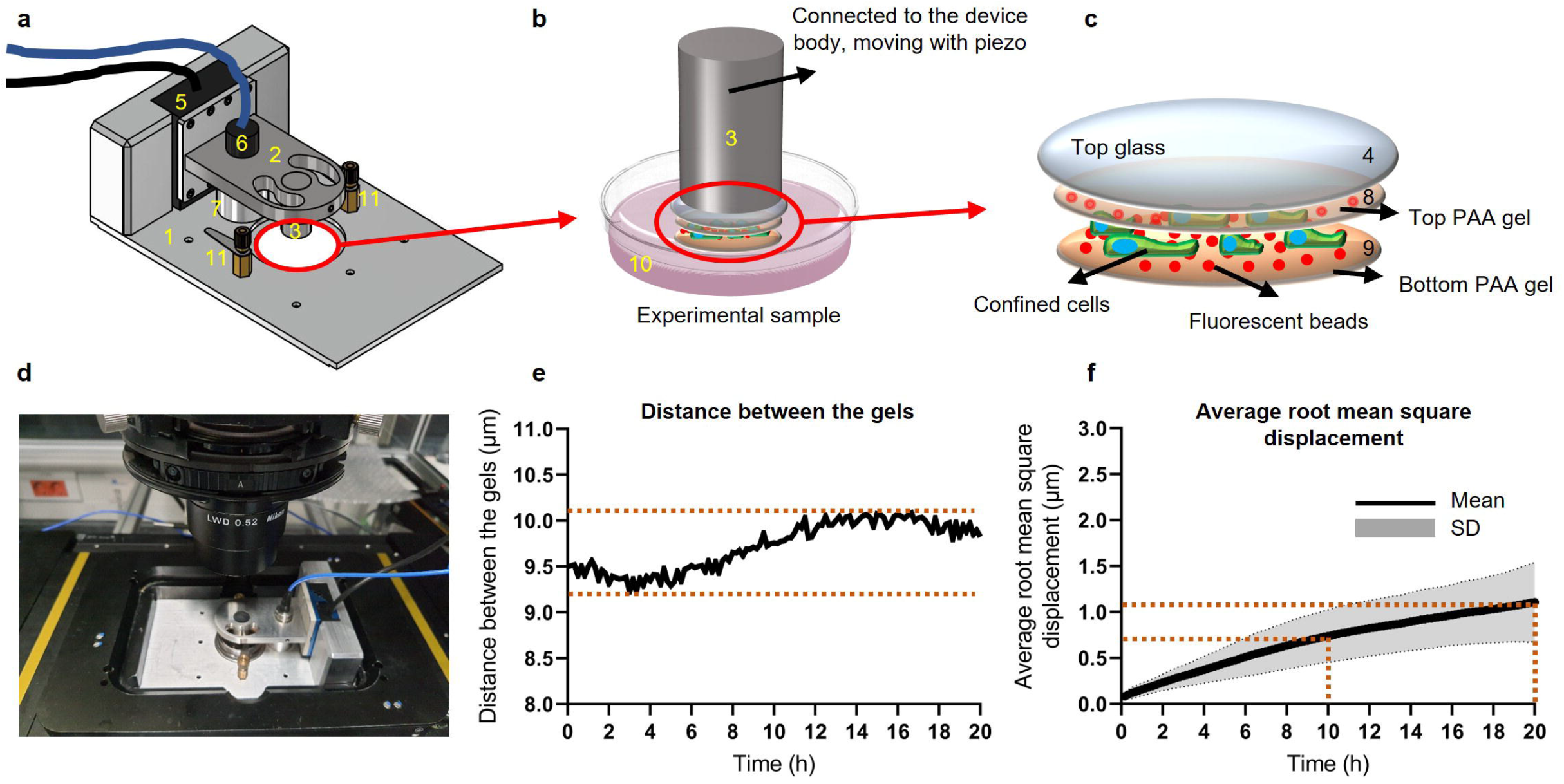
Design and characterization of CFM. (**a**) Design of CFM. The different components are designated as follows: 1. CFM body. 2. Lever arm to hold the sensor and cylinder. 3. Cylinder. 5. Piezo motor. 6. Capacitive sensor. 7. Conductive target (metal cylinder) for capacitive distance measurement. 11. Clamps to secure the Petri dish. (**b**) Schematic of the sample. 10. Glass-bottom Petri dish. (**c**) Schematic of confinement with cells seeded between two PAA gels containing fluorescent beads. 4. Glass coverslip glued to the cylinder. 8. Top PAA gel polymerized on the glass coverslip. 9. Bottom PAA gel polymerized on the glass-bottom Petri dish. (**d**) Image of CFM, mounted on a spinning-disk microscope. (**e**) An example of the confinement stability over 20 hours of imaging. The diagram shows the distance between the bottom and top gels based on time. (**f**) Root-mean-square displacement (RMSD) over 20 hours of imaging. Data are presented as mean ± SD, with the solid line indicating the mean and the shaded band indicating SD. Data represent 5 independent experiments performed on different days and on different samples. The average change in the distance between the bottom and top gels was about 700 nm after 10 hours of imaging and about 1 *µ*m after 20 hours. Illustration in panel a was created by the authors using AutoCAD, whereas the remaining panels were created by the authors using PowerPoint.

Detailed dimensions and material specifications of the individual CFM components are provided in Supplementary Note 1.

### 2.2 Confinement stability over time

One of the most important requirements for studying biological systems under confinement is to ensure a stable confinement level during a long-term experiment. In effect, the smaller the biological entity, the greater the precision and stability required of the instrument. Here, we target a low-micrometre range for the gap height, which demands high positional stability. To study and quantify the stability of the CFM, we measured the distance over a 20 h time window. Fig. 1e and Supplementary Video 1 show an example of such measurements, in which the maximum change in distance between the bottom and top gels is only about 800 nm. The stability was quantified by the average root-mean-square displacement (RMSD) of 5 experiments each conducted on a different day using a different sample (Fig. 1f). Consistent with the time series, the average drift in confinement distance over 10 h and 20 h was 700 nm and 1 *µ*m, respectively. Hence, the instrument design enables investigation of long-term effects of confinement on cellular responses, fate, and mechanical reorganization.

### 2.3 Combining traction force measurement and confinement during live imaging

The primary motivation for developing CFM was to address the lack of a method that combines tunable confinement, live-cell microscopy, and simultaneous measurement of lateral (in-plane) and normal (axial) traction stresses. After confirming confinement stability, we assessed whether CFM could vary confinement during 3D image-stack acquisition while enabling post hoc quantification of traction stresses. By embedding fluorescent beads in the PAA gels during fabrication (Fig. 1c), we tracked gel displacements under confinement and computed 3-component (2.5D) traction stresses using an adapted traction force microscopy pipeline implemented in a custom MATLAB program available on Zenodo (DOI: 10.5281/zenodo.19469501; see Methods, section “Traction force microscopy and stress calculation”).

Fig. 2a and Supplementary Video 2 illustrate stepwise confinement of a HeLa cell during live imaging, with the corresponding bottom-gel lateral (Fig. 2b) and normal stress maps (Fig. 2c). Stress maps are shown as 2D (xy) top-view heatmaps derived from the 3D stacks: “lateral” denotes in-plane traction, and “normal” denotes the axial component.

**Figure 2.**
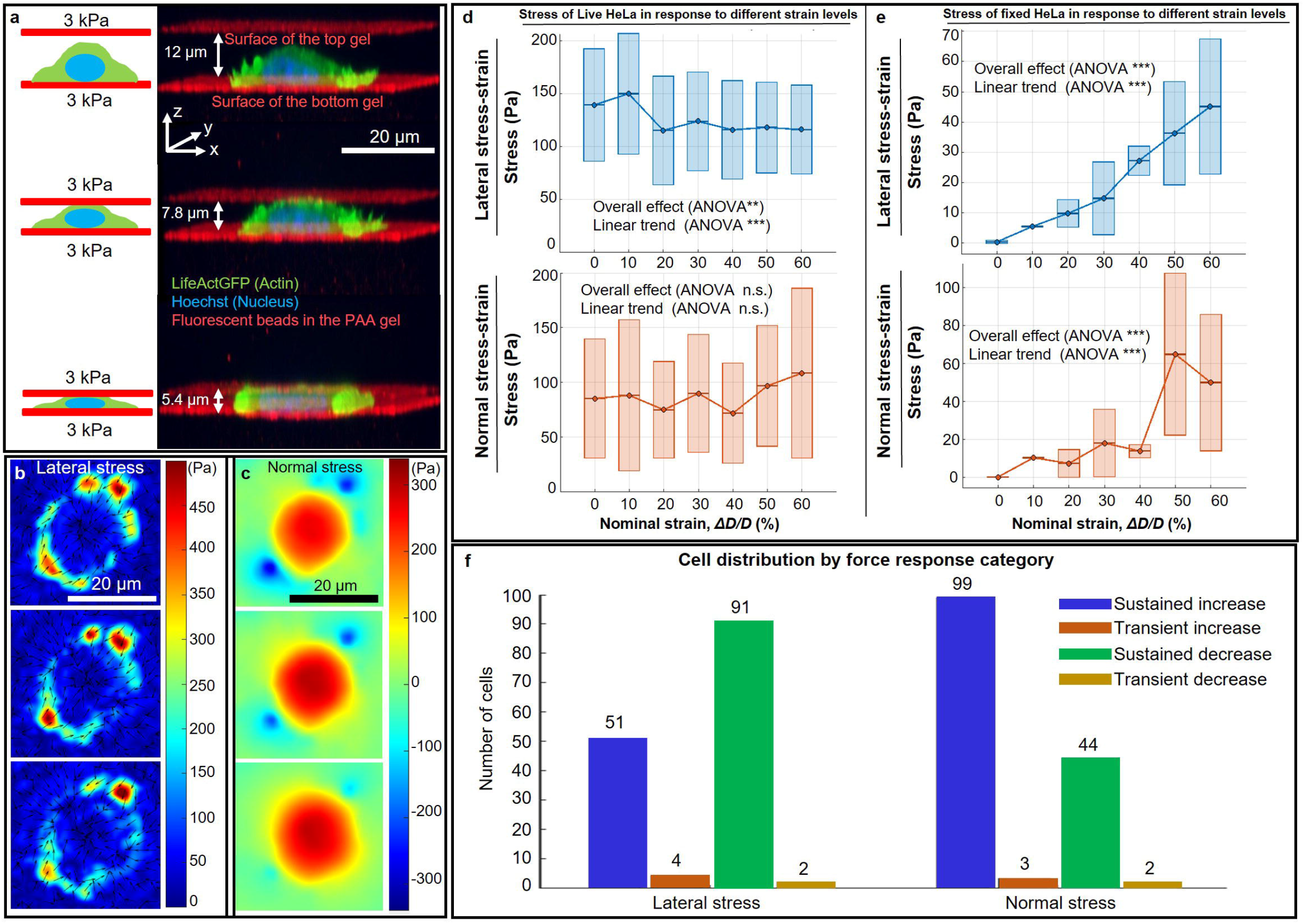
Step-by-step confinement of a HeLa cell. (**a**) 3D view of stepwise confinement during live imaging from 12 *µ*m to 5.4 *µ*m. Red planes below and above the cell indicate the bottom and top PAA gel surfaces. Gel thicknesses are 50–80 *µ*m; both gels have a stiffness of 3 kPa. (Green: Lifeact-GFP; red: fluorescent beads in the PAA gel; blue: Hoechst.) (**b**) Example lateral stress map on the bottom gel during stepwise confinement. Color bar, stress in Pa. (**c**) Corresponding z-direction stress map on the bottom gel. Color bar, stress in Pa. Positive stress indicates pushing into the substrate; negative stress indicates pulling up. Panels (**a**)–(**c**) show a representative example from the experiments in panel (**d**), which were repeated across 5 independent samples on 3 experimental days. (**d**) Stress versus nominal strain under vertical confinement of live HeLa cells. Rectangles show mean ± SD for each nominal-strain bin; the central horizontal line and circles indicate the bin mean, and lines connect bin means. Top, lateral stress: one-way ANOVA, *p* = 0.00832; Tukey–Kramer multiple comparisons, n.s.; two-sided simple linear regression, slope = −46.43, 95% CI [73.09, 19.77], *p* = 0.000692. Bottom, normal stress: one-way ANOVA, *p* = 0.157; two-sided simple linear regression, slope = 20.29, 95% CI [−9.68, 50.25], *p* = 0.184. Each observation corresponds to one confinement step from one cell and was assigned to the nearest nominal-strain bin (0.0–0.6 in 0.1 steps, using *ε*). In total, 315 observations from 105 cells were binned; unique cells contributing to bins 0.0–0.6 were 105, 21, 37, 50, 33, 33 and 26, respectively. Data were pooled from 5 independent samples across 3 experimental days. (**e**) Stress versus nominal strain under vertical confinement of paraformaldehyde-fixed HeLa cells (4% PFA). Rectangles show mean ± SD for each nominal-strain bin; the central horizontal line and circles indicate the bin mean, and lines connect bin means. Top (lateral): one-way ANOVA, *p* = 3.21 10^−15^; two-sided simple linear regression, slope = 69.23, 95% CI [58.31, 80.15], *p* = 9.62 10^−19^. Bottom (normal): one-way ANOVA, *p* = 1.31 10^−10^; two-sided simple linear regression, slope = 95.72, 95% CI [72.43, 119.01], *p* = 1.85 10^−11^. Each observation corresponds to one confinement step from one cell and was assigned to the nearest nominal-strain bin as in (**d**). Data were obtained from one sample on one experimental day; 63 observations from 21 cells were binned. Because fixed cells generate much smaller stresses than live cells, live and fixed traces are plotted on separate y-axes for readability. (**f**) Distribution of 148 cells across whole-trace lateral and normal stress-response categories, classified as sustained increase, transient increase, sustained decrease, or transient decrease, highlighting heterogeneous mechanical adaptation under confinement. Data were pooled from 10 independent samples across 9 experimental days. Stress stabilization was quantified independently from the last 15 time points (75 s) of each time series as described in Methods. For these measurements, only the gel (bead) channel was acquired, with no simultaneous cell-imaging channel.

In the example shown in Fig. 2a, the distance between the gels was reduced from ~12 *µ*m to 5.4 *µ*m. In principle, the spacing can be adjusted across the full travel range of the linear actuator (up to ~18 mm) down to 0 *µ*m, although the usable range during imaging is limited by the working distance and axial travel range of the objective. The red planes in Fig. 2a mark the bottom and top gel surfaces; PAA gel thickness is typically 50–80 *µ*m.

To further demonstrate the applicability of CFM across a range of biological length scales, we performed additional proof-of-concept experiments in diverse contexts, including immune cell migration, durotaxis under confinement, tissue-scale mechanics during Drosophila embryogenesis, cell death, and the viscoelastic characterization of multicellular cancer spheroids (Supplementary Figs. 1–5, Supplementary Videos 3–10 and Supplementary Notes 2–6).

We next examined whether traction stresses scale with confinement. We first used a multi-step assay (Fig. 2d,e) in which each cell underwent three consecutive confinement steps and stresses were quantified after each step. Confinement is reported as nominal strain, 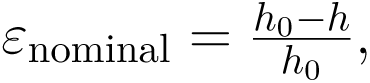 where *h*_0_ is the cell height immediately before the first confinement step and h is the height after a given step.The achieved *ε*_nominal_ was measured individually from the recorded geometry for each confinement step. Because HeLa cells vary substantially in initial height, the same absolute gel–gel spacing can correspond to different *ε*_nominal_ values; normalization therefore enables comparison across cells.

We repeated the three-step confinement protocol on 105 viable HeLa cells (∽5 min between steps) and binned strains into nominal levels (0.0–0.6 in 0.1 increments; ±0.05 tolerance). Across 105 cells, this yielded 315 stepwise observations spanning a nominal strain range of ∼5% to 60%, which were used for the analysis in Fig. 2d,e. Line segments are shown only to guide the eye and do not necessarily imply a continuous response profile for an individual cell.

As summarized in Fig. 2d, lateral stress decreased with increasing confinement (ANOVA **\*\***; linear trend **\*\*\***; pairwise Tukey’s (HSD), not significant (n.s.)), whereas normal stress showed no significant dependence on strain (ANOVA ns; trend ns). These data are consistent with active adaptation that reduces in-plane traction while preventing a systematic rise in vertical load under confinement. To verify that this behavior reflects active regulation, the experiment was repeated on fixed HeLa cells (4% PFA; 21 cells; 63 observations; Fig. 2e). In this passive condition, both lateral and normal stresses increased with strain (ANOVA **\*\*\***; trend **\*\*\***), as expected for elastic loading in the absence of active remodeling. We report stresses on the bottom gel; stresses on the top gel were not analyzed because refractive-index–induced imaging artifacts confound displacement tracking. These artifacts arise from a cell-induced lensing effect, as described previously^37^ (see Methods, section “Traction force microscopy and stress calculation”).

### 2.4 Cellular stress response to vertical confinement is heterogeneous

The multi-step confinement assay already indicated substantial cell-to-cell heterogeneity in both lateral and normal stresses at comparable strain levels, as reflected by the broad dispersion in Fig. 2d,e. To examine these differences with higher temporal resolution, we next performed a single-step confinement assay on 148 HeLa cells. In these experiments, cells were confined between two 3 kPa PAA gels, and lateral as well as normal stresses were quantified on the bottom gel. Three-dimensional imaging of the gel channel was acquired every 5 s for 100 time points. A single vertical compression step to a final confinement level of *ε*_nominal_ = 5–60% was applied shortly after time point 10 (*t* ≈ 45 s after imaging began) and completed within *<* 0.5 s. This strain range overlapped with that used in the multi-step assay (Fig. 2d,e) and enabled us to resolve how stress dynamics evolve after confinement onset.

Analysis of these single-step time series confirmed marked heterogeneity in long-term stress behavior (Fig. 2f). Some cells showed a sustained or transient increase in lateral and normal traction stresses after compression, whereas others showed a sustained or transient decrease. Nevertheless, despite this variability in both magnitude and direction, most cells reached stress stabilization by the end of imaging: 91 out of 148 cells (61.5%) for lateral stress and 109 out of 148 cells (73.6%) for normal stress. Here, “stabilized” means that over the final 15 time points (75 s), the stress trace was both flat (end slope within 3 × the median absolute deviation (MAD) of the time derivative in the tail window) and quiet (end standard deviation ≤ 0.8× the tail interquartile range). Heterogeneity quantification and stabilization criteria are described in Methods, section “Quantification of stress stabilization and trend classification”.

### 2.5 Stress dynamics correlate with morphological responses under confinement

The marked heterogeneity in stress behavior raised the question of whether these different profiles reflect underlying morphological differences or distinct cellular states. To address this, we first performed a statistical analysis relating mechanical and morphological responses under confinement using a broader subset of cells for which all variables could be scored unambiguously (*n* = 74; Supplementary Fig. 6 and Supplementary Note 7). Mechanical variables were extracted from gel-channel time series, whereas blebbing before and after imaging was scored from cell-channel images acquired immediately before and immediately after the time-lapse. This analysis revealed that a larger compression step, higher strain, and prior confinement increased the likelihood of an immediate stress jump and bleb formation, whereas blebbing was associated with a subsequent decrease in normal stress over time. These findings suggested that the apparent heterogeneity may, at least in part, reflect cells being captured at different stages of a shared adaptive response rather than exhibiting fundamentally distinct response modes. We next focused on bleb formation as a key morphological feature by repeating the single-step confinement experiments on 24 cells while imaging both the gel and cell channels continuously. We therefore used the single-step confinement protocol with high-temporal-resolution dual-channel imaging of marker beads and Lifeact-GFP, allowing precise alignment of traction-stress dynamics to confinement onset and direct comparison with morphological events such as bleb initiation and expansion. Across these experiments, we consistently observed a rapid, short-lived stress increase immediately after confinement, consistent with a passive response to abrupt compression. The magnitude of this transient force spike depended strongly on the degree of confinement. Of the 24 cells analyzed, 4 already displayed blebs at the start of imaging, 12 formed blebs during the imaging window, and 8 did not show blebbing before the end of the recording. Cells that blebbed during imaging typically showed a slower, sustained rise in traction stresses after the initial transient, indicating active adaptation to confinement. Once blebs began to expand, traction stresses stabilized or declined, marking a transition in stress dynamics associated with bleb initiation and likely pressure relief. Although bleb inflation itself is largely passive, the accompanying stress changes and redistribution likely reflect active regulation of cortical contractility, intracellular pressure, and cell–substrate adhesion.

#### 2.5.1 Transient stress increase precedes bleb formation in most cells

Temporal analysis of traction stress relative to bleb initiation revealed a clear pattern in many cells. In most cells that formed blebs under confinement within our imaging window (12 out of the 20 cells without pre-existing blebs), traction stress transiently increased shortly before and around bleb onset. In this onset-aligned subset, the stress increase was typically followed by a plateau or decline during subsequent bleb expansion (Fig. 3a–d; Supplementary Video 11). Consistently, inspection of the full dataset, including pre-blebbing or pre-confined cells that cannot be onset-aligned, showed that stresses did not continue to rise during ongoing bleb expansion but instead tended to stabilize or relax (Supplementary Video 12). These observations suggest that bleb initiation is often preceded by an active cortical contraction phase that increases intracellular pressure and transmitted traction, which is then partially relieved upon bleb expansion. Together, these results support and extend prior work showing that mechanical confinement can upregulate actomyosin activity^10,17^ and raise intracellular pressure^16^, both of which favor bleb initiation^38^; here, we directly resolve the temporal ordering, with a transient traction-stress increase preceding bleb onset in most cells. However, this trend was not universal, as some cells deviated from this pattern, highlighting intrinsic cellular heterogeneity and potentially distinct mechanical thresholds for bleb formation.

**Figure 3.**
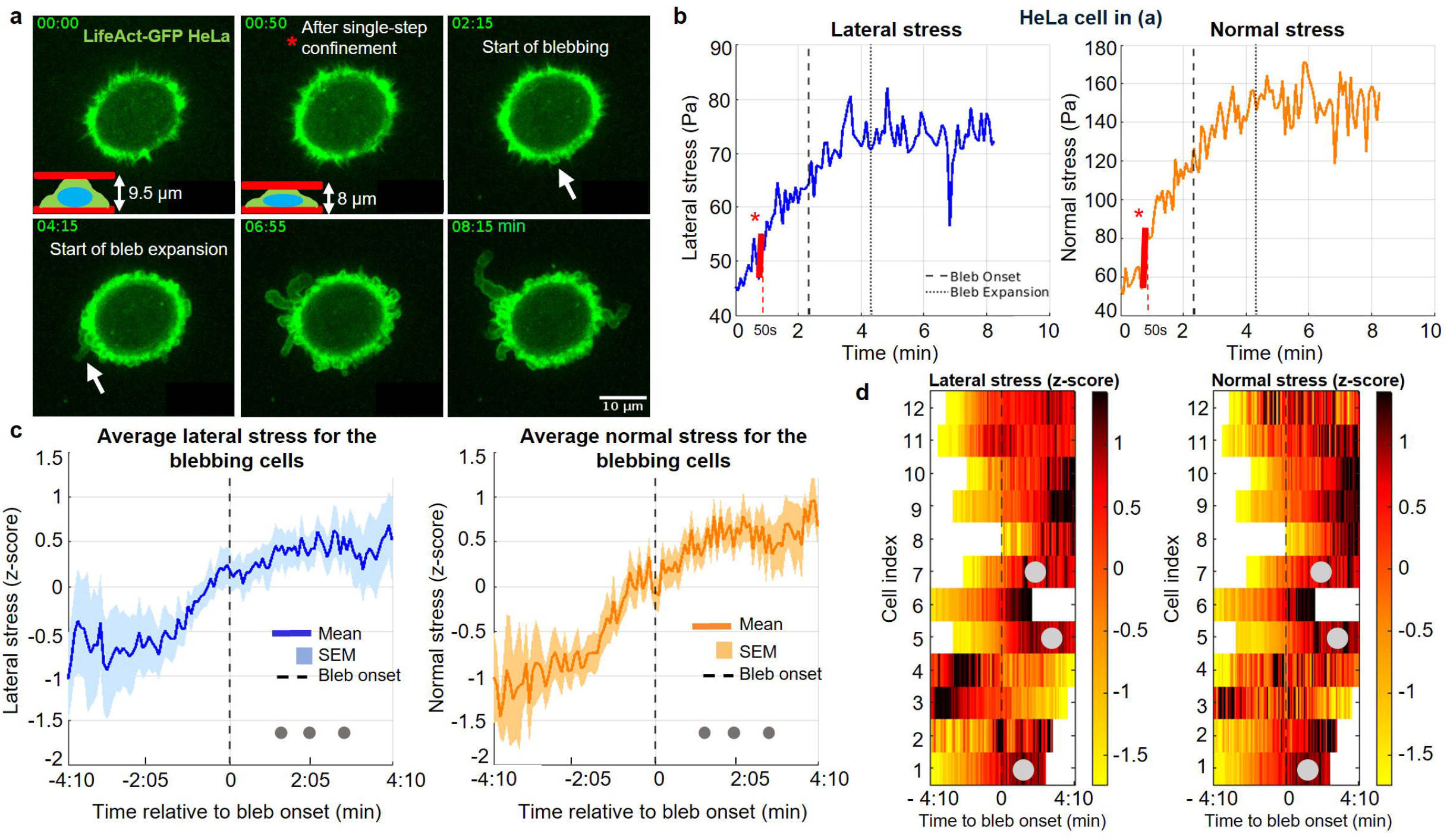
Confinement triggers a transient stress enhancement that precedes bleb onset and is followed by stabilization. (**a**) Top-view time-lapse images of a representative DMSO-treated control cell undergoing single-step vertical confinement applied between T10 (45 s) and T11 (50 s) (5 s imaging interval). Schematic overlays indicate reduction of the inter-gel spacing from 9.5 *µ*m to 8 *µ*m. White arrows indicate emerging blebs, which initiate at 2:15 (min:s), with expansion beginning at 4:15 (min:s). Scale bar, 10 *µ*m. (**b**) Lateral and normal stress on the bottom gel for the cell in (**a**) over time. A transient stress enhancement follows confinement and precedes bleb formation. The red line and star mark the immediate post-confinement stress increase. Stress stabilizes after bleb expansion begins. The pre-step rise in lateral stress likely reflects pre-existing confinement (“preconditioning”), with the cell already adapting to an earlier compression. (**c**) Averaged lateral and normal stress time series for 12 control cells from 3 independent experiments (3 samples across 3 days), shown as 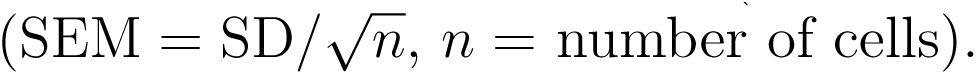 Only cells with clearly defined bleb onset during imaging were included. Dark gray dots mark bleb expansion in the 3 cells in which it occurred. Stress profiles are aligned to bleb onset (*t* = 0) and expressed as within-cell *z*-scores over the plotted window, *z*(*t*) = [*s*(*t*) *µ*_cell,window_]*/σ*_cell,window_, where *s*(*t*) is the lateral or normal stress and *µ*_cell,window_ and *σ*_cell,window_ are that cell’s mean and standard deviation over the same window, respectively. The window spans 4:10 (min:s) (50 time points at 5 s intervals) before and after bleb onset. (**d**) Heatmaps of lateral (left) and normal (right) stress *z*-scores for the 12 control cells in (**c**), aligned to bleb onset (*t* = 0) and normalized as in (**c**). Each row represents one cell and each column one time point (5 s interval); color encodes relative stress level (yellow, low; dark red, high). Alignment highlights the typical sequence across cells: stress increases before and around bleb onset, followed by a plateau or decline. Light gray circles mark bleb expansion when present, and missing or trimmed bins are shown in white. The window spans 4:10 (min:s) (50 time points) before and after onset.

#### 2.5.2 ROCK inhibition abolishes bleb formation and reduces cellular traction stresses

To further probe the basis of the observed stress dynamics, we inhibited ROCK-dependent contractility with Y-27632. Treated cells showed significantly reduced baseline traction stress compared to control cells and only a small transient stress spike under high strain upon confinement (Fig. 4a–d; Supplementary Video 13). This spike was not apparent in the signal averaged across all 16 cells. Importantly, bleb formation was completely absent in this group, even at very high strain (Fig. 4a; Supplementary Video 13), indicating that ROCK-dependent contractility is required for both the confinement-induced traction-stress response and bleb initiation under mechanical compression.

**Figure 4.**
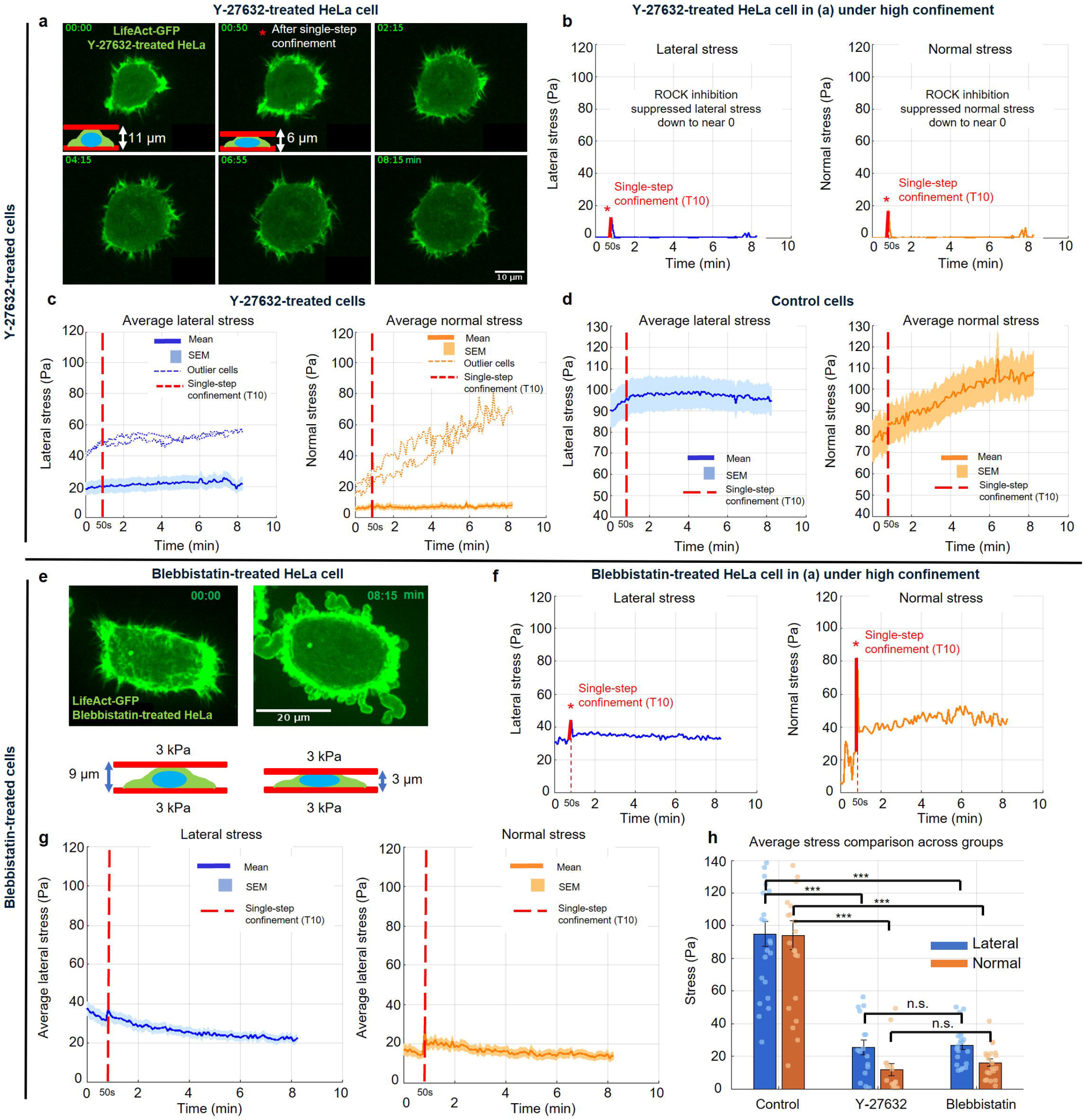
ROCK and myosin II inhibition reduce confinement-induced stress with distinct effects on blebbing. (**a**) Top-view time-lapse images of a representative Y-27632–treated cell under single-step vertical confinement applied after 00:45 min (10th time point; 5 s imaging interval). Schematic overlays show reduction of the inter-gel spacing from 11 *µ*m to 6 *µ*m upon confinement. Despite strong confinement, no bleb formation is observed in ROCK-inhibited cells. Scale bar, 10 *µ*m. (**b**) Lateral and normal stress profiles for the cell in (a) over time. Stress remains close to zero throughout, consistent with ROCK inhibition. The red line and star indicate the stress jump immediately after the confinement step. (**c**) Averaged lateral and normal stress profiles (mean SEM, where 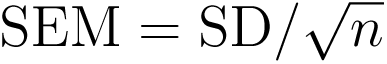 and *n* is the number of cells) for 16 Y-27632–treated cells from 3 samples across 3 independent days. Outliers were identified using the pre-defined robust criterion (median+3 MAD; union across lateral/normal); 2/16 cells (12.5%) were flagged and excluded from the mean SEM curves, but are shown as dotted overlays. (**d**) Averaged lateral and normal stress profiles (mean SEM) for 23 DMSO-treated control cells from 3 samples across 3 independent days. Stress magnitude in control cells is approximately 10-fold higher than in Y-27632–treated cells, highlighting the strong effect of ROCK inhibition on stress generation. The red dotted line in (c) and (d) marks the first time point immediately after single-step confinement. (**e**) Representative images of a blebbistatin-treated cell before and after confinement. This cell shows confinement-induced bleb formation despite blebbistatin treatment. Imaging was limited to the gel channel because of blebbistatin phototoxicity under live-cell fluorescence. Schematics below the images show reduction of the inter-gel spacing from 9 *µ*m to 3 *µ*m upon confinement. Scale bar, 20 *µ*m. Similar confinement-induced blebbing in blebbistatin-treated cells was observed across 3 independent samples collected over 3 experimental days. (**f**) Lateral and normal stress profiles over time for the blebbistatin-treated cell shown in (e), indicating stress reduction under blebbistatin treatment. (**g**) Averaged lateral and normal stress values (mean SEM, where 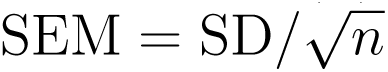 and *n* is the number of cells) from 21 blebbistatin-treated cells. Outliers were assessed using the same criterion as in (c); no cells were flagged (0/21), so mean SEM was computed over all cells. (h) Average lateral and normal stress levels (mean SEM) for DMSO control (*n* = 23 cells), Y-27632-treated (*n* = 16 cells), and blebbistatin-treated (*n* = 21 cells) groups. Each bar represents the mean stress over the full imaging period for each cell; dots indicate individual cells. Statistical comparisons were performed using one-way ANOVA followed by Tukey–Kramer multiple-comparisons testing. For lateral stress, one-way ANOVA: *p* = 1.17 10^−13^; control versus Y-27632: *p* = 3.88 10^−11^, ***; control versus blebbistatin: *p* = 5.53 10^−12^, ***; Y-27632 versus blebbistatin: *p* = 0.9900, n.s. For normal stress, one-way ANOVA: *p* = 2.27 10^−14^; control versus Y-27632: *p* = 4.62 10^−12^, ***; control versus blebbistatin: *p* = 2.30 10^−12^, ***; Y-27632 versus blebbistatin: *p* = 0.8837, n.s. Data from all groups were pooled from 3 independent samples collected across 3 experimental days.

Despite the complete absence of blebbing and the strong reduction in traction stress under confinement, some Y-27632–treated cells showed a delayed response. After extended confinement (*>* 1.5 h), a few cells developed elevated traction stress on the substrate (outliers in Fig. 4c), while blebbing remained absent throughout. (Within each sample, fields were acquired sequentially in ascending position order; thus, position index serves as a proxy for time spent under confinement, with higher indices indicating longer pre-exposure.)

#### 2.5.3 Blebbistatin reduces traction stress but does not fully prevent blebbing at high strain

Next, we tested direct myosin II inhibition using blebbistatin. To avoid light-dependent blebbistatin toxicity, we omitted continuous imaging of the cell channel during the time-lapse and acquired cell images only immediately before and after the sequence.

As expected, blebbistatin-treated cells showed reduced traction-stress magnitude relative to controls (Fig. 4e–h; Supplementary Video 14), although stress levels remained slightly higher than in the Y-27632–treated group (Fig. 4h). Blebbing was largely suppressed at low to moderate strain (*<* 50%), but at very high strain (*>* 60%), a subset of blebbistatin-treated cells still formed blebs (Fig. 4e; Supplementary Video 14).

Notably, even in blebbistatin-treated cells that blebbed, we did not observe the characteristic traction-stress dynamics seen in controls: a transient stress increase before bleb formation followed by stress stabilization or decline. Instead, stress profiles remained relatively stable over time. These findings further support that active myosin II–driven contractility is important not only for bleb initiation under physiological strain, but also for the characteristic force dynamics before and after bleb expansion. In the absence of sufficient contractile buildup, as observed after myosin II inhibition, blebs might emerge under extreme strain. We hypothesize that this likely results from alternative, passive mechanisms, such as cortical rupture induced by excessive mechanical stress.

#### 2.5.4 Contractility inhibition markedly softens cells

To track how cells relieve mechanical load after a rapid compression step, we analyzed normal-stress relaxation after confinement in control and myosin II-inhibited cells (Supplementary Fig. 7a). Because volumetric traction reconstruction requires acquisition of a full 3D gel stack, the temporal resolution was limited to ~5 s per time point. As a result, not all cells yielded relaxation profiles that could be meaningfully analyzed: in some cases, the transient stress peak and most of the subsequent decay occurred within one or two sampling intervals, rendering the response temporally undersampled and mechanically irresolvable. We therefore restricted the viscoelastic analysis to the mechanically resolvable subset of stress profiles (see Methods, section “Viscoelastic analysis of normal-stress relaxation”). As shown in Supplementary Fig. 7b, early normal-stress relaxation time constants remained in the single-digit-second range and did not differ significantly between groups (*τ*_norm_ = 7.79 ± 1.47 s, *n* = 13, control; 3.71 ± 1.29 s, *n* = 4, blebbistatin; two-sided Wilcoxon rank-sum test, *p* = 0.130). Consistent with previous reports^39,40^, the instantaneous modulus was markedly reduced by blebbistatin (*E*_0_ = 488.2 ± 63.4 Pa, *n* = 13, control; 132.1 ± 32.6 Pa, *n* = 4, blebbistatin; *p* = 0.0034) (Supplementary Fig. 7c). Effective viscosity was also lower under blebbistatin (control 567.2 ± 194.6 Pa·s, *n* = 13; blebbistatin 156.4 ± 86.2 Pa·s, *n* = 4) (Supplementary Fig. 7d). Because *η* was skewed, we analyzed log_10_(*η*); the group comparison on this scale was not statistically significant (one-way ANOVA, *p* = 0.0898). The estimated geometric-mean ratio (control/blebbistatin) was ∼3.9, with a wide 95% CI (approximately [0.39, 38.7]), consistent with reduced viscosity under blebbistatin but limited by sample size.

As discussed above, we interpret the post-step stress spike as a predominantly passive indentation response of the cell body or nucleus against the gel, transmitted through cell–ECM contacts. Under myosin II inhibition, cells are mechanically softer (Supplementary Fig. 7c) and transmit less load to the substrate, so the step-induced jump is smaller and often falls below detectability or relaxes between time points at 5 s resolution, despite no increase in baseline noise. Consistent with a passive indentation readout, blebbistatin reduces the elastic contribution to the jump without measurably altering the short early *τ* within our fixed fitting window. Thus, the “missing jump” is best explained by reduced elastic and baseline load transmission rather than by measurement noise. Blebbistatin likely also lowers pre-existing surface or cortical tension and weakens nucleus–cortex and cortex– ECM coupling, further reducing vertical load transmission. Together, these effects explain why blebbistatin-treated cells show little or no visible jump except under extreme confinement (*>* 60%; cf. Fig. 4e,f), and why *τ* provides limited contrast whereas *E*_0_ decreases robustly and *η* trends lower.

#### 2.5.5 Integrative model of bleb formation and mechanical behavior

Taken together, our results support a model in which blebbing emerges as a mechanical feedback response to actomyosin-driven cortical tension and intracellular pressure (Fig. 5). Before bleb formation, cells likely accumulate stress through contractile buildup. Upon bleb expansion, this pressure is relieved, as evidenced by the observed decrease in normal stress following blebbing events. Consistent with this, ROCK inhibition via Y-27632 treatment—which abolishes actomyosin contractility—almost completely eliminates both traction stress and bleb formation. In contrast, partial inhibition of myosin II with blebbistatin reduces stress but does not fully block blebbing, particularly at high strain levels (*>* 60%). Under such extreme confinement, blebbing can still be mechanically triggered by external forces, through cortical rupture or detachment of the membrane from the cortex, even in cells with impaired contractility.

**Figure 5.**
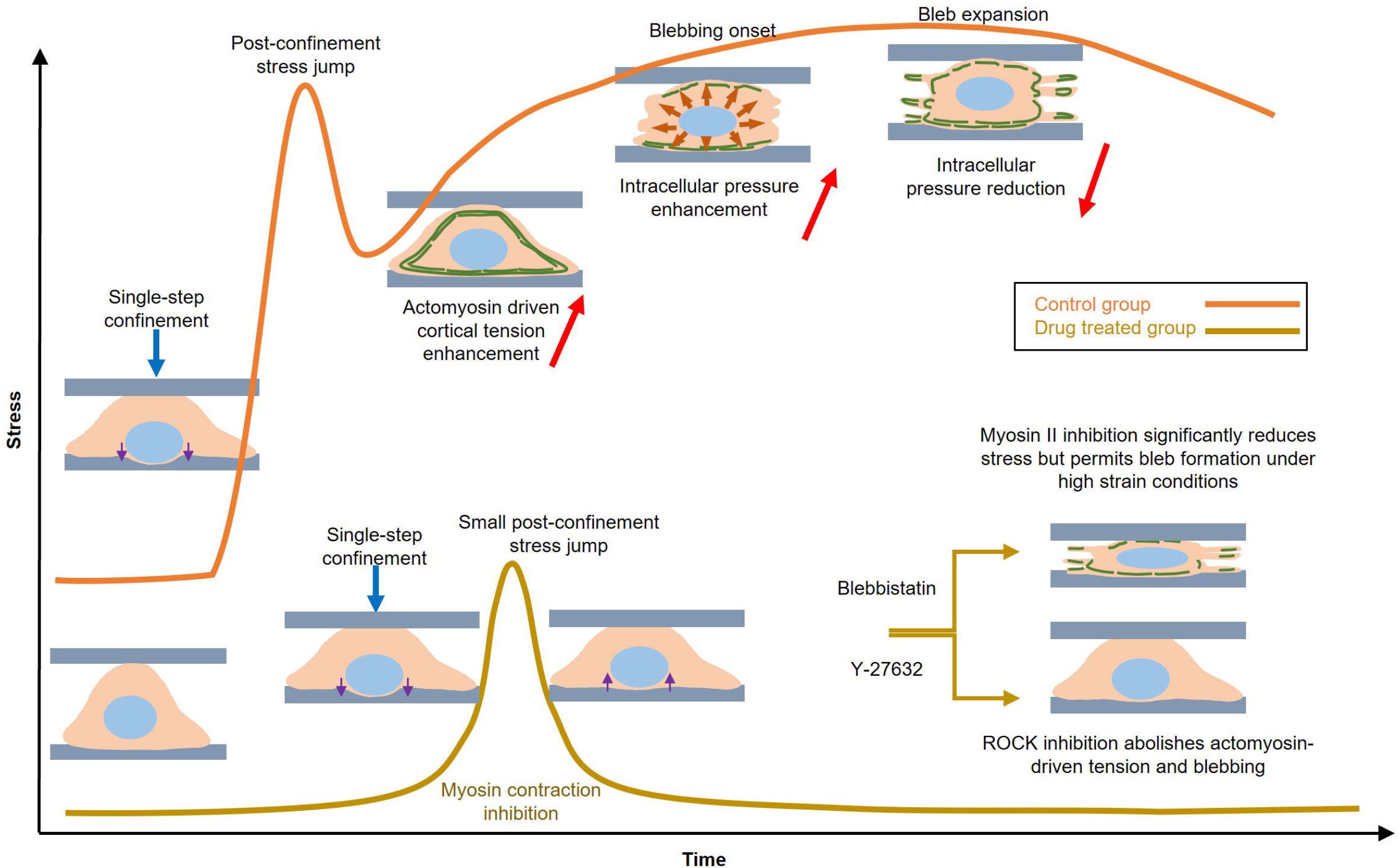
Proposed model of bleb formation and mechanical regulation under confinement. Vertical confinement increases actomyosin-dependent cortical tension, which in turn elevates intracellular pressure. This mechanical buildup can cause local membrane–cortex detachment or cortical rupture, thereby initiating bleb formation. A transient increase in traction stress precedes or accompanies bleb onset, and the newly formed bleb initially lacks a cortical actin layer. As the bleb expands and matures, intracellular pressure is likely partially relieved, and actin and myosin are recruited to rebuild the cortex; this phase is accompanied by stress stabilization or reduction. Pharmacological perturbations support this mechanism: ROCK inhibition (Y-27632) abolishes actomyosin-driven tension and blebbing, whereas myosin II inhibition (blebbistatin) markedly reduces stress and still permits bleb formation under high strain, but without the characteristic stress dynamics observed in control cells. Together, these results support a feedback model in which blebs act as a pressure-relief mechanism following cortical contraction. **Note:** The nucleus is shown schematically and is not a quantitative readout of nuclear deformation. Because no nuclear label was used in the dataset underlying this figure, in order to maintain 5 s temporal resolution and limit illumination load, nuclear shape dynamics were not measured. The nucleus is therefore depicted with a largely invariant shape across panels to avoid implying nucleus-specific conclusions. Only the final panel shows a more deformed nucleus to convey the qualitative expectation under extreme confinement. Illustrations in this figure were created by the authors in PowerPoint.

The variability observed across cells likely reflects intrinsic biological heterogeneity, including differences in cortical stiffness, cytoskeletal organization, adhesion area, and cell cycle state. These factors collectively shape how each cell balances internal force generation with external mechanical constraints, resulting in diverse stress and blebbing behaviors under compression.

Therefore, we propose the following model to explain the bleb formation process: under confinement, actomyosin and cortical contractility are upregulated via the known Rho–ROCK–myosin II signaling pathway, leading to an increase in intracellular pressure^41,42^, as measured by CFM. This elevated pressure promotes local cortical rupture or membrane–cortex detachment, thereby initiating bleb formation^43^. Subsequent bleb expansion facilitates partial release of intracellular pressure, contributing to mechanical adaptation.

### 2.6 Periodic force application to cells with arbitrary temporal and spatial patterns

Cells and tissues in the body are often subjected to periodic mechanical inputs with variable amplitudes and timescales. Bladder^44^, cartilage^45^, muscle^46^, vessels, endothelial cells^47^, cancer cells and immune cells are just a few examples. Given the importance of periodic mechanical loading, systematically characterizing cellular responses is critical for understanding the biomechanical factors that shape cell behavior^48^. Having established in single-step experiments that a sudden increase in confinement elicits a biphasic stress response with largely stable effective modulus over several minutes, we next asked whether CFM can also impose more complex, time-varying confinement patterns.

To this end, CFM enables us to apply periodic confinement with controlled spatial and temporal profiles while simultaneously tracking cell morphology and substrate stresses. As a proof of principle, we exposed HeLa cells to periodic confinement with different temporal patterns. Fig. 6a and Supplementary Video 15 show a slow oscillatory confinement of a HeLa cell with a 2.5-min period, cycling the inter-gel distance between 14.4 and 12 *µ*m. In a second example (Fig. 6b), we set the oscillation period to 42 s and programmed the feedback to produce a ±3 *µ*m sensor displacement. Here, the imaging interval was 6 s, which allowed us to capture only the bottom gel layer with a standard-speed microscope camera, but still resolved the stress-relaxation dynamics between confinement cycles. The resulting normal-stress traces on the substrate (Fig. 6b, middle plot) show that the stress amplitude remained relatively stable over the 15-min measurement, without obvious drift. Although we did not perform full viscoelastic fitting or population-level statistics for these oscillatory protocols, these examples illustrate how CFM can impose defined confinement histories and provide access to normal-stress time series under repeated loading. Using the same analysis framework as for single-step confinement, such data can be used to test whether cells gradually change their effective mechanical properties in response to frequent external compression and whether they retain a form of “mechanical memory” of past confinement. These questions are particularly relevant in contexts such as cancer cell invasion, where cells repeatedly squeeze through constrictions as they disseminate and may benefit from progressive softening to infiltrate dense tissues.

**Figure 6.**
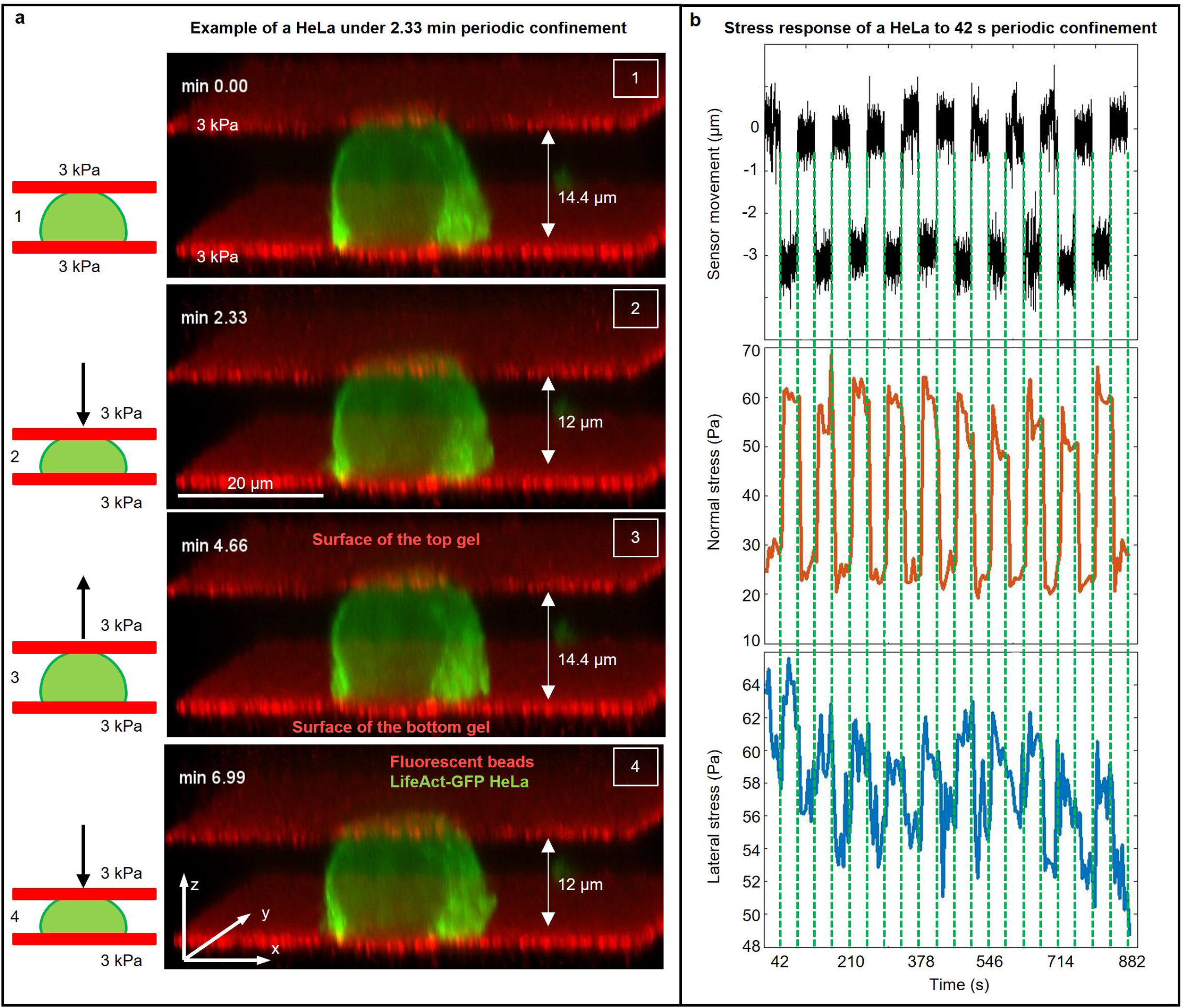
Application of periodic confinement to HeLa cells. **(a)** Representative time-lapse images showing slow, periodic confinement of a HeLa cell from 14.4 *µ*m to 12 *µ*m with a 2.5-min period during live-cell imaging. The bottom and top gels are both 3 kPa. The red planes below and above the cell show the bottom and top PAA gel surfaces, respectively. The thicknesses of the gels are 50–80 *µ*m (green: Lifeact-GFP; red: fluorescent beads in the PAA gel). **(b)** Example of rapid, periodic confinement of a HeLa cell with an oscillation period of 42 seconds. The top plot shows the sensor displacement relative to the initial position (0 = initial); the sensor moves ±3 *µ*m every 42 s. The middle plot shows the normal stress on the bottom gel; the bottom plot shows the lateral stress; both correspond to a 42 s confinement period. The inter-gel distance was 15 *µ*m at the upper sensor position and 12 *µ*m at the lower; images were acquired every 6 s. *Note:* The plots in (b) are not from the cell shown in (a). Capturing both gel surfaces every 6 s across full z-range would require high-speed volumetric imaging.

## Discussion

Cells in vivo frequently experience mechanical confinement imposed by tissue architecture, neighboring cells and the extracellular matrix^12,49^. To understand how cells adapt to such constraints we present Confinement Force Microscopy (CFM), a platform that combines dynamic vertical confinement with simultaneous measurements of lateral and normal stress components.

A central conclusion of this work is that the heterogeneous stress responses observed across confined HeLa cells are largely explained by asynchronous progression through a common adaptive pathway rather than by fundamentally distinct response types. Most cells eventually bleb after confinement, either immediately or later during the recording, indicating that blebbing is a frequent consequence of this perturbation. Aligning stress profiles to bleb onset reveals a stereotyped sequence in which substrate traction rises around bleb initiation and then relaxes as bleb expansion proceeds. Together with previous reports that confinement increases cortical contractility and intracellular pressure^10,16,17^, these findings support a model in which confinement first elevates contractility and pressure, followed by bleb formation and partial stress relief. CFM therefore provides a direct, time-resolved mechanical view of a process that has often been inferred only indirectly. The distinct effects of blebbistatin and ROCK inhibition suggest that, under extreme confinement, bleb initiation can still occur even when myosin II-dependent force production is strongly reduced, for example if confinement-induced pressure is sufficient to destabilize a mechanically weakened cortex or if other myosin isoforms partially compensate. CFM therefore reveals a partial dissociation between bleb formation and contractile force output that would be difficult to infer from morphology alone.

CFM captures both passive and active mechanical responses. The initial stress transient following confinement is reduced by contractility inhibitors, and analysis of normal-stress relaxation curves shows that these treatments soften cells. The ability to quantify force generation and mechanical material properties in the same experiment distinguishes CFM from methods that primarily report either active traction or passive rheology.

An important advantage of CFM over methods that measure only normal forces, such as parallel-plate compression^10,20,27^, microindentation or AFM-based assays^17,29^, is that it resolves force redistribution between vertical and lateral directions. The immediate response to confinement is not always identical in these two components: in some cells, normal stress increases while lateral stress transiently decreases. If only normal stress were measured, this could be interpreted as a simple increase in resistance to deformation. The lateral readout instead shows that cells can actively redistribute forces between axes. This is especially informative for blebbing: in untreated cells, bleb formation is accompanied by coordinated changes in both normal and lateral stresses, whereas in myosin II-inhibited cells, blebs can still occur despite very low forces and without a pronounced stress peak at onset. Visible blebbing should therefore not automatically be interpreted as evidence of elevated contractile force output. More broadly, simultaneous measurement of lateral and normal stresses distinguishes force-generating from force-decoupled modes of confinement adaptation.

Our data also indicate that temporal coordination between lateral and normal stresses relates to how effectively cells adapt under load. Cells with more coordinated lateral and normal stress dynamics were more likely to show a reduction in normal stress over time, whereas cells with less coordinated dynamics more often maintained or increased normal stress. This suggests that active redistribution of forces between axes is an important component of successful adaptation to confinement. In this way, CFM measures not only force magnitude but also how different stress components co-evolve in time within the same cell.

The present implementation also has several limitations. First, quantitative traction reconstruction is currently restricted to the bottom gel. Although both gels contain fluorescent beads, the upper gel is affected by imaging artefacts caused by cell-induced lensing, consistent with refractive-index-related distortions reported previously^37^. A future improvement will be the development of robust dual-surface traction readout. Second, the HeLa-cell experiments presented here focus primarily on single-step confinement, which captures one important but simplified class of mechanical perturbation. Longer recordings and more complex loading histories would be informative, but are presently constrained by fluorophore bleaching and the need to maintain imaging quality over time. Third, as with all in vitro confinement systems, the physiological relevance of CFM depends on the biological setting. Parallel-plate confinement is well suited to situations dominated by apico-basal compression, but it does not replace complementary models such as microchannels or fibrous 3D matrices, which better represent other classes of in vivo constraints.

Even with these limitations, CFM has broader potential. The proof-of-principle oscillatory experiments show that it can impose programmable confinement histories rather than only isolated steps, opening the door to studies of repeated loading, history-dependent adaptation and possible mechanical memory. More broadly, CFM should be applicable beyond the HeLa-cell paradigm studied here, including migratory immune cells, multicellular systems and cell aggregates. Rather than replacing existing confinement models, CFM adds a complementary framework in which externally imposed, time-varying compression can be linked directly to quantitative mechanical readouts. This should make it useful for dissecting how cells and tissues convert confinement into force-generating and force-relaxing behaviors across a range of mechanobiological contexts.

## Acknowledgments

F.A. thanks the GGNB Graduate School (Göttingen, Germany) and the CiM-IMPRS Graduate School (Münster, Germany). F.A. also thanks the Mechanical Workshops at the Third Institute of Physics (Göttingen) and at UKM (Münster). Figure 1a and the detailed technical drawings in Supplementary Note 1 were prepared by the mechanical workshop in Münster. The authors thank Johannes Roth (Institute of Immunology, University of Münster) for providing the HoxB8 cells and Danijela Vignjevíc (Institut Curie Paris) for providing the stably transfected HeLa cervical cancer cell lines and CT26 cancer cell lines. The authors thank Swetha Raghuraman for the cancer-spheroid protocol and Jannis Fischer for help with the supplementary videos.

## Author contributions

F.A. was involved in the design of the study, developed the instrument, designed and performed the experiments, analyzed the data, and wrote the manuscript. K.R. performed experiments on neutrophil traction stress measurements under different confinement levels. M.B. developed the traction force measurement software and adapted it to the CFM analysis. M.M. provided *Drosophila* embryos and advised on the corresponding experiments. Some of the revision experiments were performed in the laboratory of E.K., who also advised on interpretation of the results. T.B. conceived the study, developed the traction force measurement software, designed the experiments and proposed the data analysis approach. All authors discussed the results and contributed to writing the manuscript.

## Methods

### Polyacrylamide (PAA) gel fabrication

Unless otherwise stated, all chemicals were purchased from Sigma-Aldrich (Steinheim, Germany). Polyacrylamide (PAA) gels were prepared as previously described^50^, with minor modifications. In brief, for fabrication of the bottom gels, the glass-bottom dishes (CELLview 35/10 mm, Greiner Bio-One International, Kremsmünster, Austria) were washed with 0.1 N NaOH. Next, the glass was covered with 200 *µ*L of (3-aminopropyl)trimethoxysilane (APTMS) for 3 min, washed with Dulbecco’s Phosphate-Buffered Saline (PBS) 3 times and then covered with 500 *µ*L of 0.5% glutaraldehyde for 30 min. To prepare the PAA gel premix, 4 *µ*L of acrylic acid was added to 250 *µ*L of 2% N,N^r^-methylenebisacrylamide and 500 *µ*L of 40% acrylamide solution. By changing the proportion of this premix and PBS, the PAA gel stiffness was varied. For preparing the 1.2-kPa PAA gels, 75 *µ*L of the premix solution was gently mixed with 405 *µ*L of 65% PBS and 20 *µ*L of fluorescent bead solution (100 nm NH_2_-coated Micromer-RedF; Micromod, Rostock, Germany). For 9, 21, and 114 kPa gels, we mixed 113, 150, and 400 *µ*L of premix with 367, 330, and 80 *µ*L of 65% PBS, respectively. For all stiffnesses we used 20 *µ*L of fluorescent bead solution. Polymerization was induced by adding 5 *µ*L of 10% ammonium persulfate solution (APS) (Roth, Karlsruhe, Germany) and 1.5 *µ*L of N,N,N’,N’-tetramethylethylenediamine (TEMED).

For the experiments that needed cell attachment to the gel surface, we activated the acrylic-acid groups and functionalized the gel surface with a mixture of 0.2 M N-(3-dimethylaminopropyl)-N’-ethylcarbodiimide hydrochloride (EDC), 0.1 M N-hydroxysuccinimide (NHS), 0.1 M 2-(N-morpholino)ethanesulfonic acid (MES) and 0.5 M NaCl for 15 min at room temperature. Afterward, the gels were washed three times with PBS and then incubated with 60 *µ*L of 5% Bovine fibronectin (sterile-filtered, BioReagent, Merck, Germany) diluted in PBS at 37°C for 1 hour. Subsequently, gels were washed 3 times with PBS and incubated for at least another 30 min with the culture medium to be used during the live imaging. To fabricate the top gels, the same procedure was performed on microscope coverslips (12-mm round coverslips, VWR). Gel stiffness was confirmed by rheological measurements (Anton Paar Physica MCR 501, Germany) to ensure consistency.

## Cell culture

### Cell lines

HeLa and CT26 cell lines were kind gifts from Danijela Vignjevíc (Institut Curie, Paris). C2C12 cells were obtained from an established laboratory stock. HoxB8 cells were a gift from Johannes Roth (Institute of Immunology, Münster). Formal STR-based authentication was not performed in this study. Cell lines were verified by phenotype: C2C12, HeLa and CT26 by their characteristic cellular morphology and growth behavior, and HoxB8 cells by their differentiation into neutrophils. All cell lines used in this study were tested for mycoplasma contamination and were negative.

Cell lines were checked against the current ICLAC Register of Misidentified Cell Lines (version 14, released 15 February 2026). HeLa is a well-known contaminating cell line in the register; here, HeLa cells were used intentionally as an established human cervical cancer model, and their identity as HeLa was not in question. The other cell lines used in this study were not used as models of disputed identity on the basis of the information available in the ICLAC Register.

### HeLa, CT26 and C2C12 culture

HeLa and CT26 cell lines were cultured as previously described^5^. Briefly, HeLa cervical cancer cell lines and CT26 cancer cell lines were stably transduced to express Lifeact-GFP and H2B-mCherry via lentiviral transduction. All cells were confirmed negative for mycoplasma using PCR tests. HeLa, CT26, and C2C12 cells were cultured in high-glucose Dulbecco’s modified Eagle medium (DMEM) supplemented with 10% (v/v) fetal bovine serum (FBS; Capricorn, Germany) and 1% (v/v) penicillin–streptomycin (Gibco, Germany), incubated at 37 °C with 5% CO_2_ in a humidified atmosphere. The culture medium was exchanged every 2-3 days, and cells were passaged upon reaching *>* 70% confluence. To split the cells, they were first washed with PBS (modified, without calcium chloride and magnesium chloride) and suspended using trypsin–EDTA (0.05%, phenol red; Gibco). Subsequently, they were centrifuged and resuspended in the culture medium.

### HoxB8 derived neutrophil culture

HoxB8 cell lines were extracted from bone marrow of Lifeact–EGFP mice as previously described^51^ and cultivated in a clear, non-treated polystyrene 6-well plate (Falcon® 6-well Clear Flat Bottom Not Treated Cell Multiwell Culture Plate, with Lid, Sterile, Corning, USA). We used Opti-MEM 1X + GlutaMAX (Gibco, Germany) supplemented with 10% (v/v) FBS, 1% (v/v) penicillin-streptomycin solution, and 30 *µ*M *β*-mercaptoethanol (from a 50 mM stock; Gibco, Germany) as the base culture medium. Cells were suspended in the base culture medium containing 1% (v/v) SCF-conditioned supernatant (stem cell factor derived from CHO cells; a kind gift from Johannes Roth) and 10 mM *β*-estradiol. The culture medium was replaced every 2-3 days and cells were split to a density of 2 10^6^ cells per 3 mL (every second day) or to 3 10^5^ cells per 4 mL (every third day), and tested for mycoplasma contamination prior to experiments. To differentiate HoxB8 cells into neutrophils, cell suspension was centrifuged and washed twice with PBS containing 10% FBS to remove *β*-Estradiol. Subsequently, HoxB8 cells were suspended in 3 ml of base medium supplemented with 1% of SCF-supernatant in untreated six-well culture plates. Cells were differentiated for 4 days prior to the experiments.

### Cancer spheroid formation

Cell spheroids were made as previously described^5^. In brief, HeLa cervical cancer cell lines and CT26 cancer cell lines were washed with PBS (modified, without calcium chloride and magnesium chloride) and suspended using trypsin-EDTA (0.05 %), phenol red. After centrifugation they were resuspended in their culture medium. 48-well plates (Greiner Bio-One) coated with 150 *µ*L of 1% ultra-pure agarose (Invitrogen) that was cooled down for 30 min were used to prepare cancer spheroids. 1 mL of cell suspension containing 1000–2000 cells was added to each well, and the aggregates then formed spontaneously. The aggregates were collected after 2-3 days, transferred to the CFM and imaged with a 20× objective.

### Fly stocks and *Drosophila* embryo preparation

We used the *sqhAX3; sqh-Sqh-GFP* fly line (BDSC 57144) from the Bloomington *Drosophila* Stock Center. Live imaging was performed at 25 °C. Embryos were collected on apple juice agar plates smeared with yeast paste, dechorionated in bleach for 2 min, and then washed thoroughly with distilled water. Dechorionated embryos were mounted and covered with Halocarbon 700 oil (Halocarbon Products).

### Drug treatment of HeLa cells

For pharmacological inhibition of actomyosin contractility, cells were treated with either Y-27632 or blebbistatin prior to confinement experiments. Y-27632 was used at a final concentration of 20 *µ*M and incubated with cells for at least 20 min at 37^◦^C. A lower concentration of Y-27632 (10 *µ*M) was also tested and produced similar results (Supplementary Fig. 8). Blebbistatin was used at a final concentration of 50 *µ*M and incubated for 15–25 min before imaging. Blebbistatin and Y-27632 stock solutions were prepared in DMSO and protected from light throughout the procedure.

Control cells were treated with the corresponding maximum DMSO concentration (final concentration 0.1%) to account for vehicle effects. All treatments were performed in CO_2_-independent medium (Gibco, Germany) supplemented with 10% (v/v) of FBS and 1% (v/v) of penicillin–streptomycin solution immediately prior to confinement and live-cell imaging. Cells were visually monitored during treatment to confirm adherence and morphology before proceeding with mechanical assays.

### Confinement and imaging

To prepare cells for confinement and imaging, they were centrifuged and resuspended in the proper medium (differentiation medium for neutrophils and CO2-independent medium (Gibco, Germany) supplemented with 10% (v/v) of FBS and 1% (v/v) of penicillin–streptomycin solution for HeLa, CT26 and C2C12). Subsequently, they were seeded on the fibronectin-coated PAA gel at the appropriate number (20,000– 25,000 of neutrophils, 5,000–10,000 of HeLa and CT26 and 3,000–5,000 of C2C12 per gel of 12 mm diameter) and incubated for at least 30 min. Before imaging 2 ml of the proper medium supplemented with 25 mM HEPES (Millipore, Burlington, USA) was added to the petri-dish. Cancer spheroids and *Drosophila* embryos were seeded on the uncoated PAA gels. After transferring to confinement cancer spheroids were covered with the CO_2_-independent medium supplemented with 10% (v/v) of FBS, 1% (v/v) of penicillin–streptomycin solution and 25 mM HEPES while *Drosophila* embryos were covered with halocarbon 700 oil. Within the next desired time period, 4D stacks were captured at the required time intervals (based on the experiment) and desired z plane distance using the spinning disk system (CSU-W1 Yokogawa) combined with a heating chamber set at 37°C for the cells and spheroids and 25°C for *Drosophila* embryos. In order to investigate the effect of confinement, the glass bottom petri-dish containing the bottom PAA gel and seeded cells was placed in CFM. The position of the bottom gel was recognized using the fluorescent microscopy and subsequently the distance between the bottom and the top gels was adjusted, based on the question and the size of the entity in between, by moving the top gel downward. Cells were imaged with a 60 × objective and cancer spheroids and *Drosophila* embryos were imaged using a 20 × objective, utilizing the Slidebook software (3I, USA) and analyzed with ImageJ.

### Traction force microscopy and stress calculation

After imaging the confined cells for the desired time duration and with the appropriate time interval, the upper layer was moved upwards and 100 *µ*L of 10% sodium dodecyl sulfate (SDS) as a membrane lysis buffer was gently added. Afterward, the upper layer was moved back to the previous position and the relaxed bottom and top gels were captured as the reference (i.e. null force) images. Traction stresses in X-Y (Lateral) and Z directions (Normal) for single cells were calculated as previously described^50,52^. In short, PAA gel deformations were calculated based on the displacements of the fluorescent beads on the gel surface with respect to the relaxed gel after the SDS addition. Displacements were calculated by an iterative free-form deformation algorithm using the open-source software Elastix^53^. To analyze with the Elastix method, using the cubic B-spline functions, the 3D reference image was iteratively deformed and compared to the respective time point of the 3D image of the deformed gel. The iteration process started with distributing the control points of the B-spline functions over a regular mesh. The control points’ positions and, therefore, the applied deformation were adjusted in each iteration to optimize the quality of agreement between the two images measured by the advanced Mattes mutual information metric. To do so, the tuning steps followed an adaptive stochastic gradient descent algorithm^52^. The calculation progressed in a three-level pyramid approach from the coarsest to the finest scale and the comparison between images in each iteration was not carried out on the full images but on a randomly sampled subset of each image. For embryo and spheroid experiments the displacement calculation was performed differently. The lateral displacement was derived using a PIV algorithm that compared image sub-windows by an upsampled cross-correlation employing a discrete Fourier transform (DFT) method^50,54^, which uses the phase difference between the DFTs of two images to achieve subpixel resolution. The z-displacement in this approach was determined by a subpixel registration of the PAA gel surface as introduced recently^50^. The final deformation result was inserted into the equation for the Tikhonov-regularized inverted elasticity problem for finite-thickness substrates. We used a Bayesian estimate of the regularization parameter that includes a background estimate obtained by explicitly marking cell-free regions, following the approach introduced by Huang et al^55^. The equation was solved in the Fourier domain^56,57^ using a custom-made MATLAB (MathWorks) program yielding the 3-component (2.5D) traction stress exerted on the gel’s surface^52^.

The stresses reported here are the average over the cell–substrate contact area on the bottom gel, computed using a cell mask generated in ImageJ. Taken together, this analysis yields the full vectorial traction-stress components (*σ_x_, σ_y_, σ_z_*) at the cell–substrate interface of the bottom gel. In line with current nomenclature in the field, we therefore refer to our approach as a 3-component (2.5D) traction force reconstruction: all three components of the traction vector are obtained from 3D bead displacements, but they are resolved on a single planar substrate rather than throughout a surrounding 3D matrix. We do not attempt volumetric stress reconstruction around the entire cell body in the present implementation.

Finally, we note that although fluorescent beads were embedded in both the bottom and top PAA gels, we restrict quantitative traction reconstruction to the bottom interface in this study. In control experiments, traction inversion on the top gel yielded substantial apparent bead displacements and reconstructed stresses even when the top gel was positioned tens of micrometres above the cell (i.e. without physical contact), indicating non-physical signals consistent with refractive-index– induced imaging artefacts in the upper gel^37^. In principle, such artefacts can be reduced by refractive-index–matching additives (e.g. iodixanol-based media such as OptiPrep), but under our experimental conditions these additives altered cell adhesion and morphology and thereby changed the mechanical readouts, so we did not use them in this study. We therefore report only bottom-gel stresses throughout and interpret the net loading using force-balance arguments.

## Statistical analysis and visualization

3D renderings were generated using an ImageJ 3D script. Plots were generated with Python, MATLAB, and GraphPad Prism. Each figure caption lists the statistical tests applied. Statistical comparison between two groups was performed depending on whether the data were classified as normally distributed at the 5% level. Box plots display the first to the third quartile of the data, and the centre line represents the median value. The whiskers extend from the edges of the box to the furthest data point in the 1.5 × interquartile range.

For plots showing population-averaged stress time series (mean±SEM), we excluded rare traces with abnormally large stress levels to prevent a small number of extreme cells from dominating the mean. Outliers were identified automatically using a fixed, pre-defined robust threshold applied to each cell’s typical stress level over the full recording: for each cell we computed the temporal median of its lateral (or normal) stress trace, and flagged the cell as an outlier if this value exceeded the population median plus *k* times the median absolute deviation (MAD; scaled to a Gaussian-equivalent standard deviation by 1.4826), with *k* = 3:

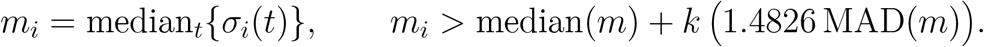

To ensure consistency between components, we used a common exclusion set defined as the union of cells flagged by the lateral and/or normal criteria; the same cells were therefore excluded from both averaged traces. Excluded traces are still shown in the corresponding plots as dotted overlays, and only the mean±SEM curves were computed after outlier removal. This outlier procedure was applied only to population-averaged time-series plots and not to summary statistics/bar plots unless explicitly stated in the figure caption.

For the viscoelastic comparison shown in Supplementary Fig. 7, group differences between control and blebbistatin-treated cells were assessed using two-sided Wilcoxon rank-sum tests.

### Correlation matrix analysis

To investigate statistical associations between mechanical and morphological variables under confinement, we computed a Pearson correlation matrix based on single-cell metrics derived from force and imaging analyses. A dataset comprising multiple features—such as blebbing behavior, confinement time, applied strain, lateral and normal stress dynamics, and per-cell stress correlations—was compiled into a structured Excel file for analysis.

The correlation matrix was calculated in Python (version 3.11) using the pandas, Seaborn and Matplotlib libraries. The dataset was loaded from an Excel file with the openpyxl engine, and pairwise Pearson correlation coefficients were computed in pandas. The resulting correlation matrix was visualized as a heatmap using Seaborn, with annotations, a color scale ranging from 1 to +1, and gridlines to facilitate interpretation. Pearson correlation was used to quantify linear associations across the dataset.

### Quantification of stress stabilization and trend classification

To determine whether traction stresses had reached a stable level toward the end of the recording, we analyzed the last 15 time points (~75 s at 5 s per time frame) of each stress time series. The traces were first smoothed with a 7-point moving average, and a linear regression was fit to this tail segment as a function of time. We then calculated the temporal derivative within this window and used its median absolute deviation (MAD) as a robust estimate of the local noise level. A cell was classified as “stabilized” if (i) the absolute slope of the tail was smaller than three times the MAD of the derivative and (ii) the standard deviation of the tail segment did not exceed 0.8 times its interquartile range. Traces that did not fulfill these criteria were classified as non-stabilized. Traces with fewer than five valid time points were excluded from the analysis.

To describe the overall behavior of each stress trace across the full recording, we further classified the smoothed full-length time series according to its baseline level, end level and maximal excursions. The baseline was defined as the mean of the first five time points and the end level as the mean of the last five time points. Upward and downward excursions were calculated as the maximum and minimum deviation from baseline, respectively, and the end change was defined as the difference between the end level and the baseline. To determine whether these changes exceeded the noise level of the trace, we estimated a robust amplitude threshold from the point-to-point fluctuations of the smoothed signal as

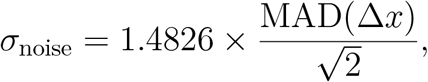

where Δ*x* denotes the frame-to-frame difference of the smoothed trace, and defined the classification threshold as *A*_thr_ = 3 ×*σ*_noise_.

Based on these quantities, each trace was classified as *sustained increase* if the end level remained above baseline by more than *A*_thr_, and as *sustained decrease* if the end level remained below baseline by more than *A*_thr_. Traces were classified as *transient increase* or *transient decrease* if they showed a clear upward or downward excursion beyond *A*_thr_ during the recording but returned to within ± *A*_thr_ of the baseline by the end. Traces that did not meet any of these criteria would have been classified as showing *no clear change*; however, no cells in our dataset fell into this category.

### Viscoelastic analysis of normal-stress relaxation

To quantify stress relaxation after a rapid confinement step, we fit a single-exponential decay to the normal-stress time series (Supplementary Fig. 7a). Because volumetric traction reconstruction requires acquisition of a full 3D gel stack, the temporal resolution was limited to ~5 s per time point. We therefore restricted the viscoelastic analysis to traces with relaxation dynamics slow enough to be resolved at this sampling rate. Because this criterion preferentially excludes the fastest, lowest-amplitude relaxations, which occur more frequently under myosin II inhibition, it cannot artificially produce a softening effect. If anything, the reported *E*_0_ and *η* values are conservative, and the true blebbistatin-induced softening is likely at least as strong.

To ensure a sufficient number of post-peak points for fitting, we applied a conservative dynamics-based pre-filter: the three time points immediately following the peak had to be lower than the peak value. The peak was defined as the first observed maximum in the stress trace after a single-step confinement, usually the first post-step time point, *t*_0_. This criterion does not impose a mechanistic model, but excludes cases in which the true peak and subsequent relaxation occurred between sampling intervals and therefore could not be temporally resolved.

Fits were performed from the peak onward using a short fixed window (typically 5–6 time points) for all included cells (5 s imaging interval) and were constrained to monotonic decays (positive amplitude and *τ*) using a robust loss function. The fit model was

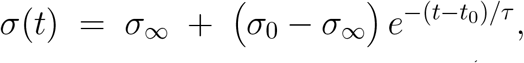

where *σ*(*t*) is the normal stress, *t*_0_ is the peak time (the first sample after the confinement step), *σ*_0_ = *σ*(*t*^+^) is the immediate post-step stress, *σ* is the apparent plateau within the fit window, and *τ* is the relaxation time constant.

From the normal-channel fits, we computed viscoelastic parameters via a Standard-linear-solid (SLS) mapping:

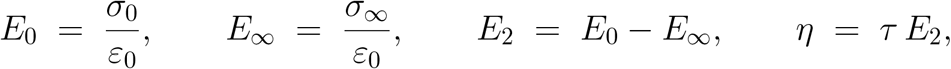

where *ε*_0_ is the measured compressive strain for the step, calculated from cell heights before and after single-step confinement, *h*_0_ and *h*_1_, as

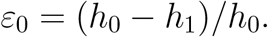

Here, *E*_0_ is the instantaneous modulus, *E _∞_*is the late modulus within the fit window, *E*_2_ is the Maxwell spring constant, and *η* is the effective viscosity.

## Data Availability

The minimum dataset necessary to interpret, verify and extend the findings of this study has been made publicly available via Figshare at https://doi.org/10.6084/m9.figshare.30051319. Source data underlying the figures are provided with this paper where applicable. Additional data are available from the corresponding author upon reasonable request.

## Code Availability

The custom MATLAB code used for traction-force reconstruction and stress analysis is publicly available on Zenodo at DOI: 10.5281/zenodo.19469501. The custom LabVIEW program used to control the CFM instrument is publicly available on Zenodo at DOI: 10.5281/zenodo.19535276.

## Statistics & Reproducibility

No statistical method was used to predetermine sample size. To demonstrate the method, we used sample sizes sufficient to assess reproducibility, with cells typically measured from at least three independent samples collected on three different days. Sample sizes were guided by the initial variance and the observed data distribution. The exact sample size (*n*) for each experiment is indicated in the corresponding figure legends.

Data were excluded if the cellular phenotype raised concerns about viability. Data were also excluded if the analysis software failed to load or process the files, if imaging was interrupted and resulted in discontinuities, or if the analysis pipeline did not return a valid output.

The experiments were not randomized. The investigators were not blinded to allocation during experiments and outcome assessment.

All experiments were independently repeated as indicated in the corresponding figure legends. Representative images are shown for experiments that yielded similar results across independent repeats. Statistical tests used for each analysis are specified in the corresponding figure legends and Methods sections. Exact *p* values are reported where possible.

