## Supplementary Information for "CFM: Confinement Force Microscopy—a dynamic, precise and stable microconfiner for traction force microscopy in spatial confinement"

Fatemeh Abbasi *et al.*

### Contents

Supplementary Figure 1. Effect of confinement level and microenvironmental stiffness on neutrophils and their nuclei S2
Supplementary Figure 2. Neutrophil undergoing 3D durotaxis S3
Supplementary Figure 3. Measuring stress during *Drosophila* embryo cellularization S4
Supplementary Figure 4. Measuring forces during cell death S5
Supplementary Figure 5. Measuring viscoelasticity of cancer spheroids S6
Supplementary Figure 6. Pearson correlation heatmap of cellular mechanical responses, blebbing behavior, and confinement parameters S7
Supplementary Figure 7. Myosin II inhibition markedly reduces stiffness ($E_0$) with a trend toward lower viscosity ($\eta$) S8
Supplementary Figure 8. Y-27632 at 10 $\mu$M reproduces the suppression of blebbing and traction stresses under single-step confinement S9
Supplementary Table 1. Method comparison for studying cells under confinement with force readouts S10
Supplementary Videos S12
Supplementary Note 1. Technical design, dimensions and material specifications of the CFM device S15
Supplementary Note 2. Effect of confinement level and microenvironment stiffness on neutrophil migration and its nucleus S24
Supplementary Note 3. Neutrophil 3D durotaxis S26
Supplementary Note 4. Measuring force generation during development S27
Supplementary Note 5. Measuring stresses imposed on the microenvironment during cell death S28
Supplementary Note 6. Measuring the viscoelastic properties of cancer cell spheroids S29
Supplementary Note 7. Statistical association between mechanical and morphological responses under confinement S30
Supplementary References S31

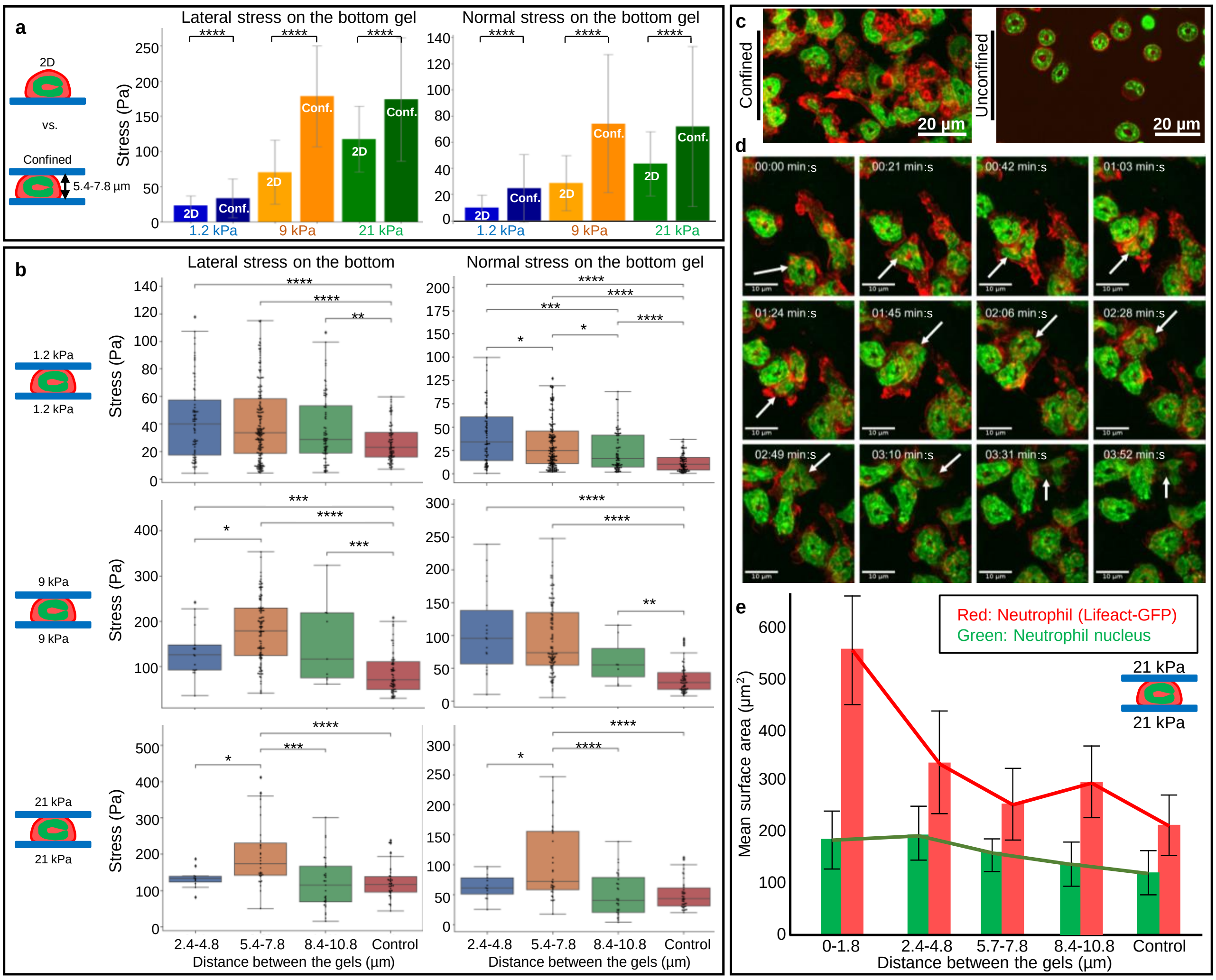


**Supplementary Figure 1:** Effect of confinement level and microenvironmental stiffness on neutrophils and their nuclei. (**a**) Median lateral and normal traction stresses on the bottom gel for unconfined neutrophils on one-layer gels (2D) and confined neutrophils in two-layer gel sandwiches at 1.2, 9 and 21 kPa. In the confined condition, neutrophils were positioned between two gels of the same stiffness at an inter-gel distance of 5.4–7.8 $\mu$m. Error bars indicate SD. Individual data points are not overlaid in (**a**) because the same confined-cell measurements are shown again in greater detail in (**b**) after subdivision by confinement range, which improves figure readability. Statistical significance in (**a**) and (**b**) was assessed using pairwise post hoc t-tests: ****, $p < 0.0001$; ***, $p < 0.001$; **, $p < 0.01$; *, $p < 0.05$; n.s., $p \geq 0.05$. Exact $p$ values are provided in the source data. Data represent $n = 177$ cells from 5 experimental days for 1.2 kPa confined, $n = 109$ cells from 9 experimental days for 9 kPa confined, $n = 33$ cells from 6 experimental days for 21 kPa confined, and $n = 83$, 68 and 41 cells from 4, 3 and 3 experimental days for the corresponding 2D conditions. (**b**) Median lateral and normal traction stresses on the bottom gel for confined neutrophils at three inter-gel distance ranges (2.4–4.8, 5.4–7.8 and 8.4–10.8 $\mu$m) and three stiffnesses (1.2, 9 and 21 kPa). The control condition had an inter-gel distance of $\sim$30 $\mu$m to prevent contact with the top gel. (**c**) Representative top-view images of unconfined neutrophils and neutrophils confined at 4.2 $\mu$m between gels with bottom and top stiffnesses of 1.2 and 21 kPa, respectively, showing pronounced nuclear shape changes under confinement (red, Neutrophil-LifeactGFP; green, nucleus stained with SYTO™ Red Fluorescent Nucleic Acid Stain Sampler Kit, No. 60 for confined cells and Hoechst for unconfined cells). Scale bar, 20 $\mu$m. (**d**) Representative time-lapse images of a confined neutrophil undergoing marked nuclear deformation while encountering another cell along its migration path under the same gel conditions as in (**c**) (red, Neutrophil-LifeactGFP; green, nucleus stained with SYTO™ Red Fluorescent Nucleic Acid Stain Sampler Kit, No. 60). Scale bar, 20 $\mu$m. Panels (**c**) and (**d**) are representative of experiments repeated under the indicated conditions with similar results. (**e**) Mean cell and nucleus surface area under the three confinement ranges and the control condition on 21 kPa gels. Error bars indicate SD. Data represent $n = 200$ cells from 3 experimental days.

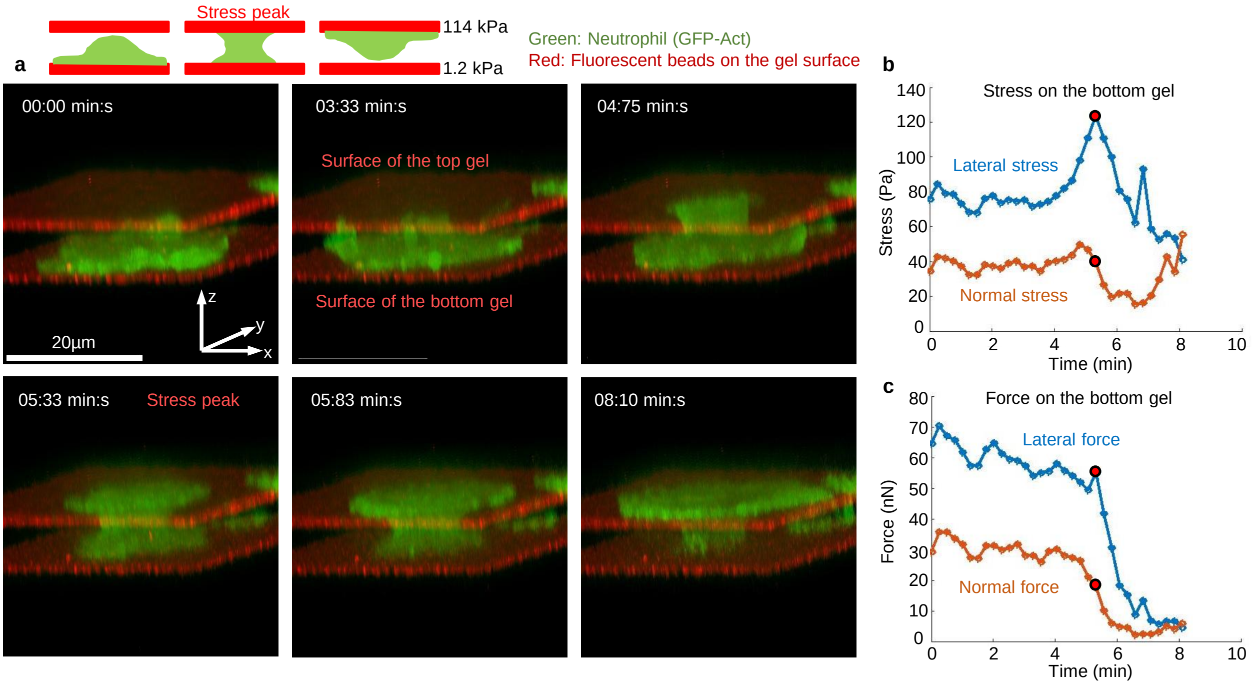


**Supplementary Figure 2:** Neutrophil undergoing 3D durotaxis. (**a**) Representative time-lapse images of a neutrophil undergoing 3D durotaxis. The bottom PAA gel is 1.2 kPa and the top gel is 114 kPa. Initially, the neutrophil is seeded on the soft bottom gel and spreads; over the next 8 min 10 s it migrates to the stiff top PAA gel, begins to spread on it, and detaches from the soft bottom gel. The red planes below and above the cell represent the bottom and top PAA gel surfaces, respectively. Gel thicknesses were typically 50–80 $\mu$m. (Green: neutrophil actin, Lifeact–GFP; red: fluorescent beads in the gel.) (**b**) Lateral and normal stresses on the soft bottom gel during durotaxis. Red dots mark the times of the lateral-stress peak, which correspond to the cell's cylindrical shape between the gels during transfer (5:33 min:s in the images in panel a). (**c**) Lateral and normal forces on the soft bottom gel during durotaxis. Red dots indicate the time of the lateral-stress peak, after which the lateral force on the soft bottom gel drops sharply as the neutrophil spreads fully on the stiff top gel and detaches from the soft bottom gel.

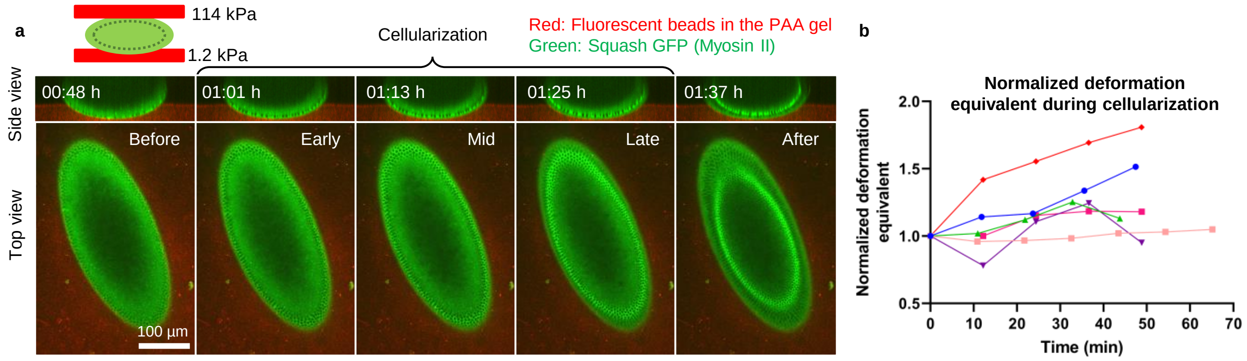


**Supplementary Figure 3:** Measuring stress during *Drosophila* embryo cellularization. (**a**) Side and top views of a representative *Drosophila* embryo expressing fluorescent Squash-GFP (green) during cellularization. The red channel represents the fluorescent labels of the PAA gels. The five time points depict the cellularization stages (time is measured from the start of imaging). Scale bar, 100 $\mu$m. Panel (**a**) is representative of experiments on $n = 6$ embryos imaged across 3 independent days with similar results. (**b**) Normalized equivalent deformation during cellularization of *Drosophila* embryos. Data are from $n = 6$ embryos imaged across 3 independent days.

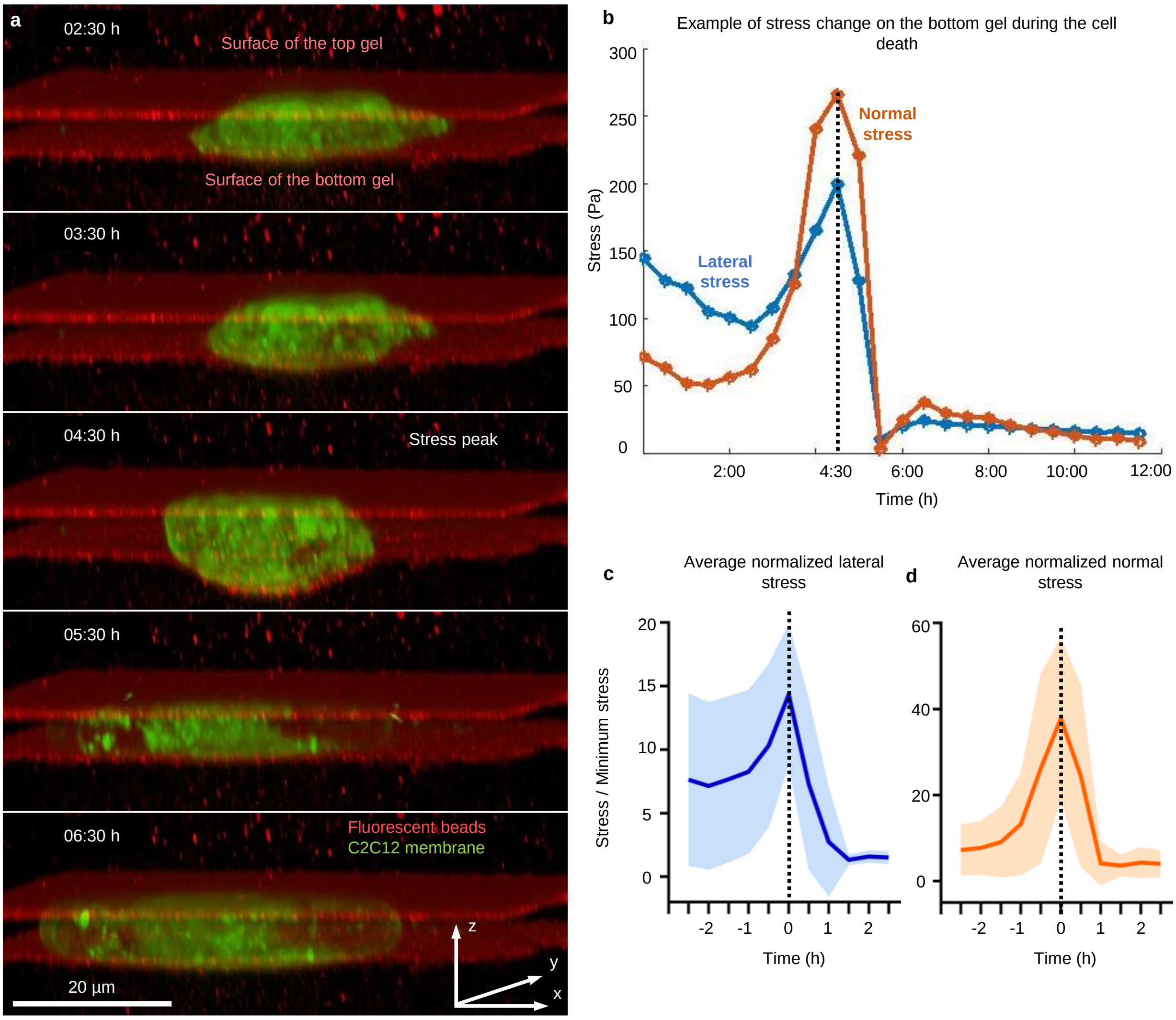


**Supplementary Figure 4:** Measuring forces during cell death. (**a**) Representative images of a C2C12 cell confined between two fibronectin-coated PAA gels (soft bottom: 1.2 kPa; stiff top: 21 kPa) over 12 h, during which the cell undergoes cell death. Scale bar, 20 $\mu$m. Panel (**a**) is representative of experiments on $n = 8$ cells from one sample with similar results. (**b**) Time courses of lateral and normal stresses on the bottom gel for the cell in (**a**); a stress peak occurs at 4:30 h after the start of imaging. (**c**) Average lateral stress on the bottom gel 2.5 h before and 2.5 h after the cell-volume increase (black dashed line). Data represent $n = 8$ cells from one sample. (**d**) Average normal stress on the bottom gel for $n = 8$ cells 2.5 h before and 2.5 h after the cell-volume increase (black dashed line). *Note:* C2C12 cells were labeled with CellMask™ Deep Red plasma membrane stain.

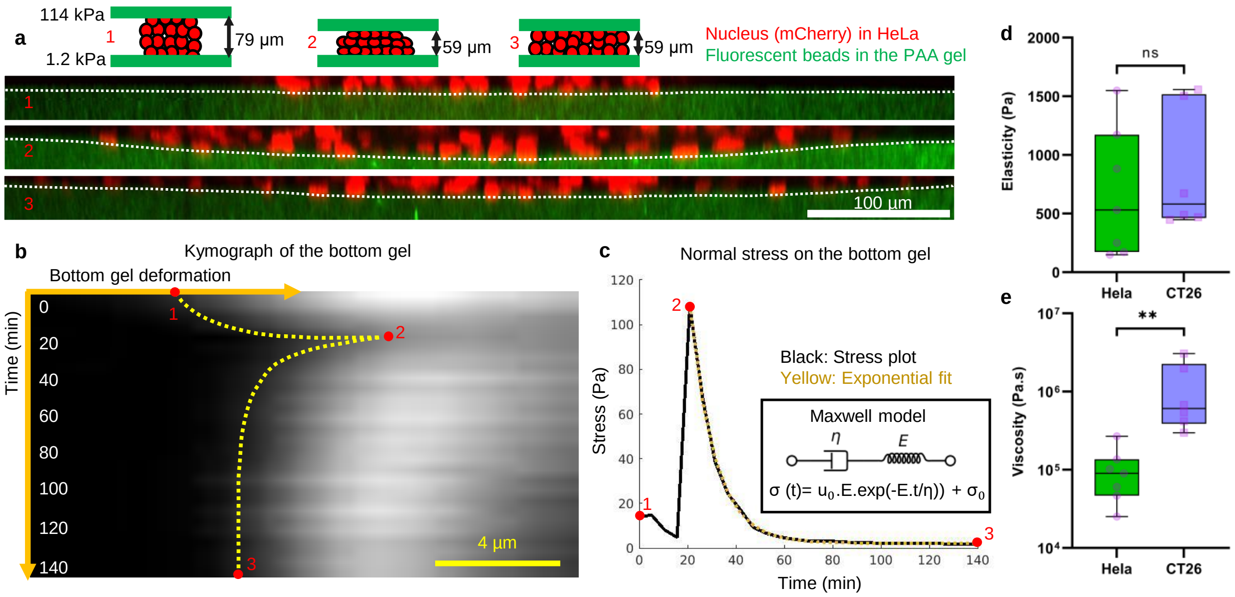


**Supplementary Figure 5:** Measuring viscoelasticity of cancer spheroids. (**a**) Representative time-lapse images of a HeLa spheroid confined between two uncoated PAA gels—a soft bottom gel (1.2 kPa) and a stiff top gel (114 kPa). Label 1 shows the beginning of the experiment in which the spheroid has minimal contact with the top gel. Label 2 shows the moment when the spheroid is compressed between the gels and the deformation and stress on the bottom gel reach a maximum. Label 3 shows the end of the experiment in which the inter-gel distance is the same as in label 2, but the stress has relaxed over time. Scale bar, 100 $\mu$m. Panel (**a**) is representative of experiments on $n = 7$ HeLa spheroids performed across 3 independent days with similar results. (**b**) Kymograph of bottom-gel deformation over time. Scale bar, 4 $\mu$m. (**c**) Normal stress on the bottom gel over time. Labels on the plot correspond to those in panel (**a**). The yellow dotted curve is an exponential fit (Maxwell model). (**d**) Median elastic modulus $E$ for HeLa and CT26 spheroids. Data represent $n = 7$ HeLa spheroids and $n = 6$ CT26 spheroids measured across 3 independent days. Box plot: box = first to third quartile, center line = median, whiskers = furthest data point within 1.5×IQR; Kolmogorov–Smirnov test, $p > 0.1$. (**e**) Median viscosity $\eta$ for HeLa and CT26 spheroids. Data represent $n = 7$ HeLa spheroids and $n = 6$ CT26 spheroids measured across 3 independent days. Box plot: box = first to third quartile, center line = median, whiskers = furthest data point within 1.5×IQR; Mann–Whitney test, $p = 0.0012$.

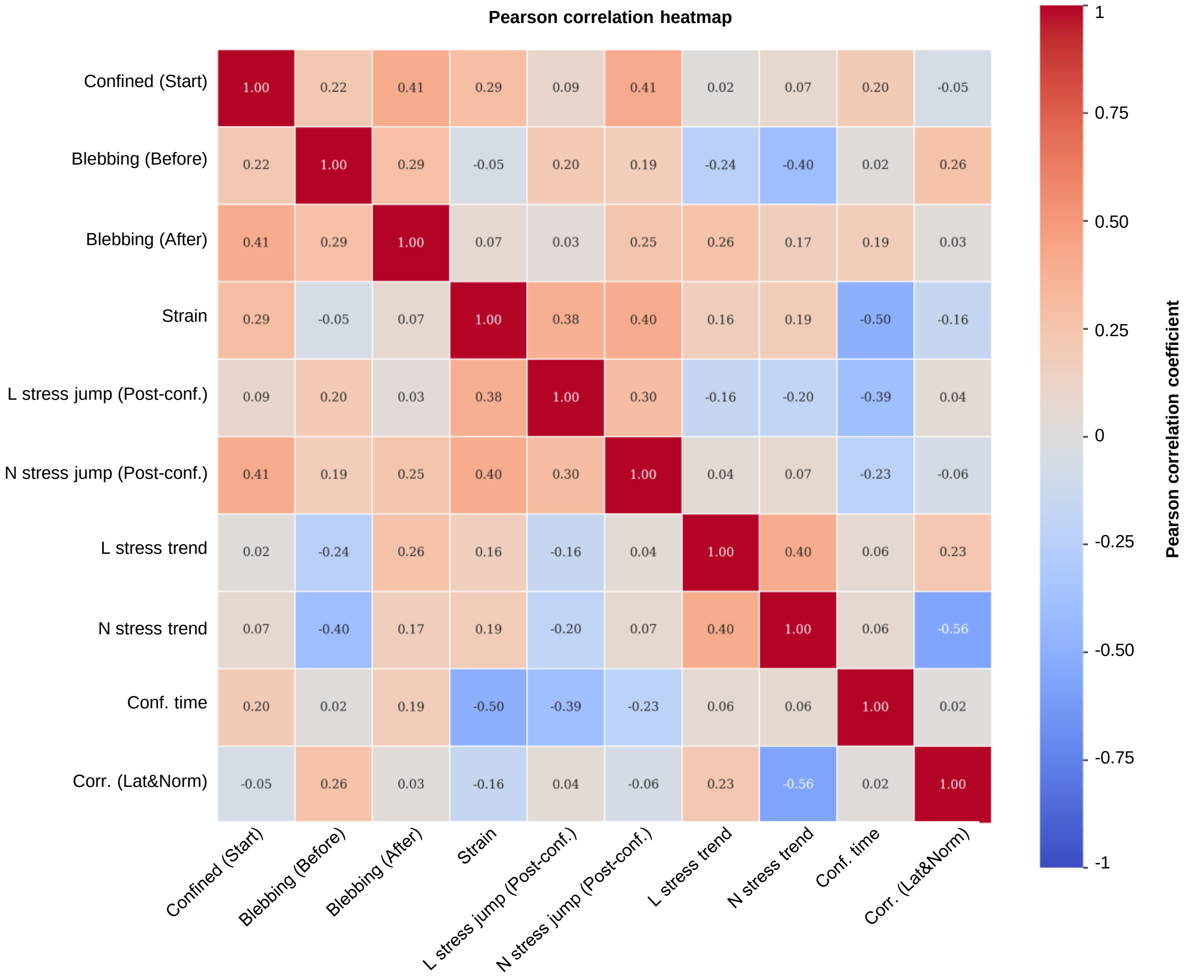


**Supplementary Figure 6: Pearson correlation heatmap of cellular mechanical responses, blebbing behavior, and confinement parameters.** The heatmap displays the Pearson correlation coefficients among key mechanical and morphological variables analyzed across confined HeLa cells: *Confined (Start)* — whether the cell was already under confinement at the start of imaging; *Blebbing (Before)* — whether the cell exhibited blebs before imaging began; *Blebbing (After)* — whether blebs were present in the cell image acquired immediately after the time-lapse; *Strain* — the relative change in cell height, calculated as the difference between pre- and post-confinement height normalized to the initial height; *L stress jump (Post-conf.)* — whether a detectable lateral-stress jump occurred after the single-step compression; *N stress jump (Post-conf.)* — whether a detectable normal-stress jump occurred after the single-step compression; *L stress trend* — the direction of lateral stress over time (increase, decrease, or no clear change); *N stress trend* — the direction of normal stress over time (increase, decrease, or no clear change); *Conf. time* — proxy for elapsed time under confinement, represented by imaging position index within each sample; *Corr. (*Lat & Norm*)* — the Pearson correlation coefficient between lateral and normal stress over the imaging period. Strong positive correlations highlight linked mechanical and morphological transitions, whereas negative correlations suggest decoupling between variables under compression. Strain correlated positively with both lateral ($r = 0.38$) and normal ($r = 0.40$) stress jumps, consistent with stronger immediate mechanical responses at higher compression. Confinement time showed negative correlations with strain ($r = -0.50$) and lateral stress trend ($r = -0.56$), suggesting time-dependent mechanical adaptation. Correlation between lateral and normal stress dynamics was positively related to confinement time ($r = 0.45$). Only the strongest correlations are described; all values reflect Pearson's $r$. Variables were coded as follows: yes $= 1$ and no $= 0$ for *Confined (Start)*, *Blebbing (Before)*, *Blebbing (After)*, *L stress jump (Post-conf.)*, and *N stress jump (Post-conf.)*; *L stress trend* and *N stress trend*: increase $= 1$, no clear change $= 0$, decrease $= -1$; *Strain*, *Conf. time*, and *Corr. (*Lat & Norm*)*: continuous variables. Data are from $n = 74$ cells pooled from 10 independent samples collected across 9 experimental days.

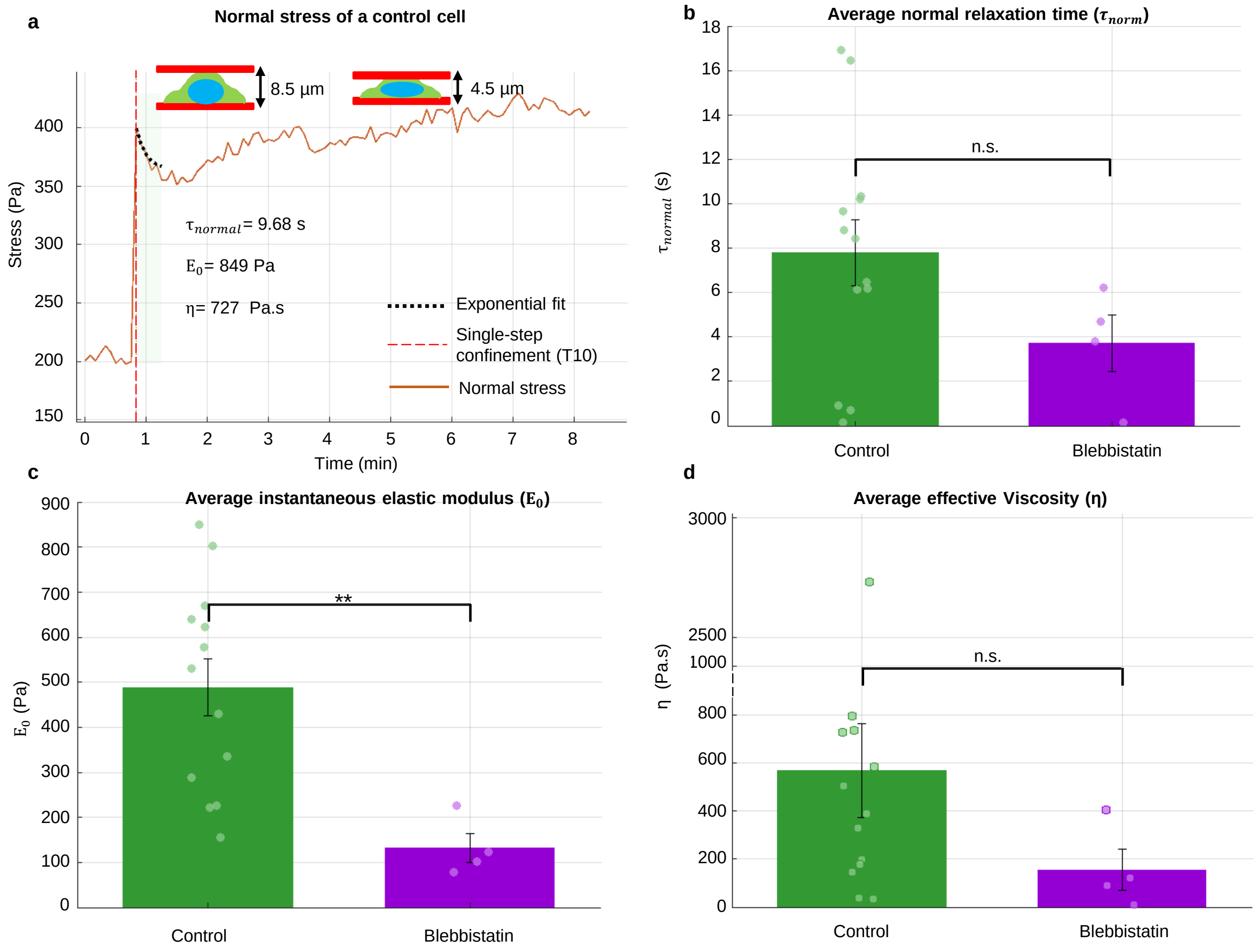


**Supplementary Figure 7: Myosin II inhibition markedly reduces stiffness ($\boldsymbol{E_0}$) with a trend toward lower viscosity ($\boldsymbol{\eta}$).** (**a**) Example normal (axial) stress trace from a control cell with single-exponential fit, $y(t) = C + Ae^{-(t-t_0)/\tau}$, fitted over a fixed window of 5–6 time points after the post-step peak (5 s per time frame). For this cell, the viscoelastic readouts were $E_0 = 849$ Pa, $\eta = 727$ Pa·s, and $\tau = 9.68$ s. (**b**) Group comparison of normal relaxation time $\tau_{\mathrm{norm}}$; bars show mean values, error bars indicate SEM, and dots indicate individual cells. Control, $n = 13$; blebbistatin, $n = 4$; two-sided Wilcoxon rank-sum test, $p = 0.130$. (**c**) Instantaneous modulus $E_0$ in the normal channel; bars show mean values, error bars indicate SEM, and dots indicate individual cells. Control, $n = 13$; blebbistatin, $n = 4$; two-sided Wilcoxon rank-sum test, $p = 0.00336$. (**d**) Effective viscosity $\eta$ in the normal channel; bars show arithmetic mean values, error bars indicate SEM, and dots indicate individual cells. Control, $n = 13$; blebbistatin, $n = 4$. Because $\eta$ was skewed, statistical comparison was performed on $\log_{10}(\eta)$ using one-way ANOVA, $p = 0.0898$. The estimated geometric-mean ratio between control and blebbistatin was approximately 3.9, with a wide 95% confidence interval of approximately [0.39, 38.7], consistent with a trend toward lower viscosity under blebbistatin but limited by sample size. Cells were included only if the three samples immediately following the peak were strictly lower than the peak, and all fits used the same short window of 5–6 time points after the peak. Here we report viscoelastic parameters only for the subset of cells whose post-step relaxation was slow enough to be resolved at 5 s per time frame.

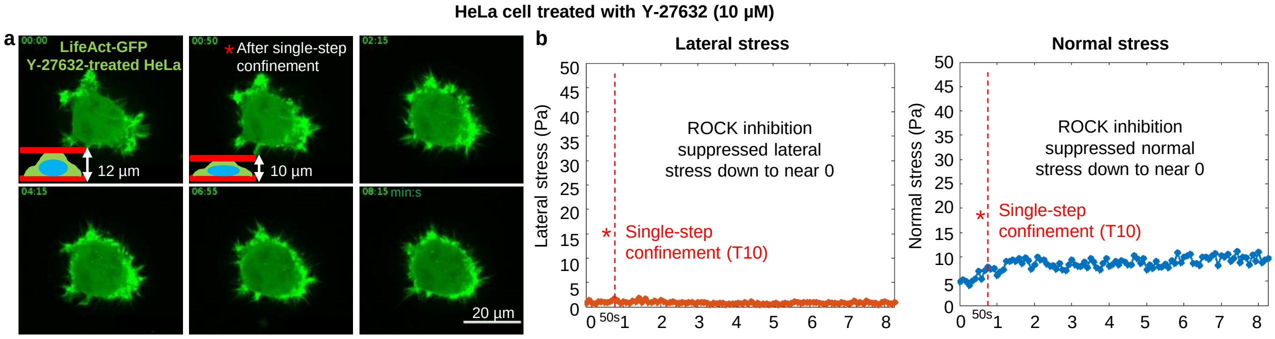


**Supplementary Figure 8: Y-27632 at 10 $\mu$M reproduces the suppression of blebbing and traction stresses under single-step confinement.** **(a)** Representative time-lapse of a HeLa LifeAct-GFP cell during a single-step confinement from 12 $\mu$m to 10 $\mu$m (nominal strain $\sim$16%), showing no bleb formation throughout the recording (scale bar, 20 $\mu$m). Consistent with this example, no blebbing was observed across analyzed cells, including experiments at much stronger confinement (down to 2–3 $\mu$m). **(b)** Lateral (blue) and normal (orange) traction-stress time series for the same cell, remaining near zero and within the noise level over the full imaging window. In untreated controls, comparable confinement steps elicit substantially larger stress magnitudes and a detectable post-step stress increase, confirming that 10 $\mu$M Y-27632 is sufficient to strongly suppress force generation and the associated mechanical response.

| **Platform** | Dynamic confinement | Dual-axis traction (lat.&norm.) | Spatial map | Live-cell imaging | Typical throughput | Output (force/stress) | **Notes / Key (dis)advantages** |
|---|---|---|---|---|---|---|---|
| **Confinement Force Microscopy (CFM)** | ✓ | ✓ | ✓ | ✓ | **High**† | **$\tau_{lat}$ map, $\tau_{norm}$ map (time-resolved)** | **Dynamic gap control with simultaneous lateral *and* normal stress maps (dual-axis); post hoc 3D bead-based reconstruction; high-NA compatible.** |
| AFM indentation / parallel-plate compression [1–3] | ✓ | ✗ | ∼point/line | ✓ | Low | $F_{norm}$, $\delta$; apparent $E$ (global) | Normal force/indentation only; no stress map; precise control, low spatial coverage. |
| Parallel planes with spacers (fixed gap) [4–7] | ✗ | ✗ | ✗ | ✓ | High | $F_{norm}$ (if instrumented) otherwise none | Very stable fixed height; gap not tunable during imaging; typically no stress readout. |
| MicroSquisher/MicroTester (micro-compression) [8] | ✓ | ✗ | ✗ | ✓ (side view) | Low–Medium | $F_{norm}$; $\sigma = F/A$ (global) | Controlled compression + normal force; side-view limits high-NA/subcellular imaging; no stress mapping. |
| Cytocompression (parallel-plate) [1] | ✓ | ✗ | ✗ | ✓ (side view) | Low | $F_{norm}$; $\sigma = F/A$ (global, no map) | Uniaxial compression with force–displacement/recovery readouts; no stress map; single-cell, low throughput. |
| Microfluidic channels (fixed geometry) [9–14] | ✗ | ✗ | ✗ | ✓ | High | none by default (add-ons possible[13,14]) | Fixed height; not tunable during imaging; high reproducibility and throughput; generally no stress mapping at the substrate. |
| 2D Traction Force Microscopy (planar gel) [15,16] | ✗ | ✗ lateral only | ✓ | ✓ | High | $\tau_{lat}$ map | Lateral stress map only; no axial/normal component; no dynamic vertical confinement. |
| Micropost (pillar) arrays [17,18] | ✗ | ✗ lateral only | ✓ discrete | ✓ | Medium | lateral forces at posts (discrete field) | Discrete lateral forces at pillar tips; axial/normal forces typically inaccessible; fixed geometry. |
| 3D TFM in ECM gels (collagen/PA) [19] | ✗ no gap control | ✗ assumptions | ✓ | ✓ | Medium | 3D stress map | Full 3D stress in compliant matrices; no controlled vertical confinement; not a solid–interface normal stress. |

**Supplementary Table 1: Method comparison for studying cells under confinement with force readouts.** *Legend:* ✓ = present; ✗ = not available; ∼ = approximate/limited. *Abbreviations:* lat = lateral; norm = normal; NA = numerical aperture; $F_{norm}$ = normal force; $\delta$ = indentation depth; $E$ = apparent elastic modulus; $\sigma = F/A$ = average stress; $\tau_{lat}, \tau_{norm}$ = lateral/normal traction stress; "map" = spatial field; "discrete" = per-pillar values.

† "High" for CFM assumes field tiling (multiple cells/position), short–moderate time series, and batched reconstruction; the 3D TFM analysis step is the rate limiter. Throughput labels are qualitative.

*Supplementary Table 1 (continued)*

| Platform | Confinement geometry (which directions are constrained) | Typical in vivo situations best represented by each geometry |
|---|---|---|
| **Confinement Force Microscopy (CFM)** | **1D plate-like apico–basal compression between parallel plates (vertical constraint).** | **Externally imposed apico–basal compression against an extended substrate (e.g. crowded epithelial/endothelial layers; cells beneath stiff capsules/bone; compressed tissue slabs/organoids).** |
| AFM indentation / parallel-plate compression [1–3] | Local-to-global normal loading via point or plate indentation (constrains primarily the vertical direction; a true 1D parallel-plate confinement only in plate configurations). | Controlled normal loading/indentation mechanics (cell stiffness/viscoelasticity); confinement relevance depends on probe/plate geometry and whether a true parallel-plate configuration is used. |
| Parallel planes with spacers (fixed gap) [4–7] | 1D plate-like apico–basal confinement at fixed height (constant gap between parallel plates). | Stable, sustained confinement at a well-defined height (plate-like tissue compression/crowding) when dynamic loading histories are not required. |
| MicroSquisher/MicroTester (micro-compression) [8] | 1D uniaxial compression between surfaces with global loading (primarily vertical constraint; side-view imaging). | Externally imposed compression with global force readout; useful for compression/recovery mechanics; confinement relevance depends on probe/plate geometry and whether a true parallel-plate configuration is used. |
| Cytocompression (parallel-plate) [1] | 1D uniaxial/parallel-plate compression with global normal-force readout (primarily vertical constraint; side-view imaging). | Single-cell compression/recovery assays and plate-like compression scenarios dominated by normal loading. |
| Microfluidic channels (fixed geometry) [9–14] | 2D cross-sectional confinement in microchannels (restricts both width and height; pore-/capillary-like constrictions). | Cell transit through constrictions (e.g. leukocyte extravasation, immune-cell squeezing through narrow passages, cancer-cell passage through vessels/pores). |
| 2D Traction Force Microscopy (planar gel) [15, 16] | No confinement (planar 2D interface); no imposed geometric constraints beyond adhesion to a flat substrate. | 2D spreading/migration on compliant substrates; adhesion and traction dynamics without spatial restriction. |
| Micropost (pillar) arrays [17, 18] | No confinement (planar 2D interface with discrete elastic supports); no imposed geometric constraints beyond adhesion to posts. | 2D migration/spreading with per-adhesion force readout on engineered compliant substrates; not intended to model confinement. |
| 3D TFM in ECM gels (collagen/PA) [19] | 3D confinement within an ECM network (embedding in a deformable, often fibrous matrix that constrains the cell in all directions). | Cells embedded in interstitial ECM (e.g. mesenchymal migration through collagen-rich stroma; endothelial invasion into ECM) where fiber mechanics and 3D constraints dominate. |

### Supplementary Videos

#### Supplementary Video 1

**3D visualization of the top and bottom PAA gels positioned at an 8 µm inter-gel spacing.** The video demonstrates the mechanical stability of the confinement over a 20-hour imaging period. Fluorescent beads embedded in the top and bottom gels are shown in red, serving as fiducial markers to visualize and monitor gel alignment throughout the experiment. Scale bar: 20 µm.

#### Supplementary Video 2

**3D visualization of stepwise cell confinement during live imaging, from 12 µm down to 5.4 µm.** Confinement steps are applied at 5-minute intervals. The red planes above and below the cell represent the surfaces of the top and bottom PAA gels, respectively. Each gel has a thickness of 50–80 µm and a stiffness of 3 kPa. Fluorescent markers indicate cellular and gel components: green denotes F-actin (LifeAct-GFP), red marks fluorescent beads embedded in the PAA gels, and blue labels the nucleus (Hoechst). Scale bar: 20 µm.

#### Supplementary Video 3

**Example of confined neutrophils at an inter-gel distance of 3.6 µm between the bottom and top polyacrylamide (PAA) gels.** The bottom gel has a stiffness of 1.2 kPa, while the top gel is 21 kPa. The second and third parts of the video display the lateral and normal stress, respectively, exerted by the confined neutrophils on the bottom gel. F-actin, labeled with LifeAct-GFP, is shown in white in the grayscale images. Scale bar: 20 µm.

#### Supplementary Video 4

**Representative side and top view videos of confined neutrophils at an inter-gel spacing of 4.2 µm between the polyacrylamide (PAA) gels.** The bottom and top gels have stiffnesses of 1.2 kPa and 21 kPa, respectively. The video highlights a pronounced change in nuclear morphology under confinement (blue). Actin in neutrophils is labeled with LifeAct-GFP (red), and nuclei are stained with the SYTO™ Red Fluorescent Nucleic Acid Stain Sampler Kit, Nr. 60 (green). Scale bar: 50 µm.

#### Supplementary Video 5

**Representative top-view video of unconfined neutrophils seeded on a polyacrylamide (PAA) gel with a stiffness of 9 kPa.** The video highlights the characteristic doughnut-like morphology of the multi-lobular neutrophil nucleus. Actin is labeled with LifeAct-GFP (red), and the nucleus is stained with Hoechst (green). Scale bar: 50 µm.

#### Supplementary Video 6

**Representative video of a neutrophil undergoing 3D durotaxis, along with lateral and normal stress maps on the bottom polyacrylamide (PAA) gel.** The bottom and top PAA gels have stiffnesses of 1.2 kPa and 114 kPa, respectively. Initially, the neutrophil is seeded and spread on the soft bottom gel. Over the course of 8 minutes and 10 seconds, it migrates toward the stiffer top gel, begins to spread on it, and eventually detaches from the bottom gel. The red planes below and above the cell represent the surfaces of the bottom and top PAA gels, respectively. Each gel has a thickness of 50–80 µm. Neutrophil actin is labeled with LifeAct-GFP (green), and fluorescent beads embedded in the PAA gel are shown in red. Scale bar: 20 µm.

**Supplementary Video 7**

**Representative video of a neutrophil undergoing 3D durotaxis from the soft top gel to the stiff bottom gel.** The top and bottom polyacrylamide (PAA) gels have Young's moduli of 1.2 kPa and 114 kPa, respectively. The neutrophil is initially seeded and spread on the soft top gel, then migrates toward the stiffer bottom gel, begins to spread there, and ultimately detaches from the top gel. Red planes mark the surfaces of the bottom and top PAA gels. Each gel is 50–80 µm thick. Neutrophil F-actin (LifeAct–GFP) is shown in green; fluorescent beads embedded in the PAA are shown in red. Scale bar: 20,µm.

**Supplementary Video 8**

**Representative video of *Drosophila* cellularization**, showing top and side views of an embryo expressing fluorescent Squash-GFP (green). The red channel marks the fluorescently labeled PAA gels. Time in the video is indicated relative to the start of imaging. Scale bar: 100 µm.

**Supplementary Video 9**

**Representative video of a C2C12 cell confined between a soft bottom (1.2 kPa) and a stiff top (21 kPa) fibronectin-coated polyacrylamide (PAA) gel for 12 hours, undergoing cell death.** The cell membrane is labeled with CellMask Deep Red plasma membrane stain (green), and fluorescent beads embedded in the PAA gels are shown in red. Scale bar: 20 µm.

**Supplementary Video 10**

**Representative top- and side-view videos of HeLa and CT26 spheroids positioned between two uncoated polyacrylamide (PAA) gels—a soft bottom (1.2 kPa) and a stiff top (114 kPa) gel.** At a defined time point, the top gel is lowered to apply confinement, after which the inter-gel distance is maintained to monitor stress transmission onto the bottom gel. HeLa spheroids exhibit a more fluid-like behavior, with cells actively invading out of the spheroid. In contrast, CT26 spheroids show higher surface tension, with superficial cells remaining strongly adherent to the spheroid body. Scale bar: 200 µm.

**Supplementary Video 11**

**Time-lapse sequence of a DMSO-treated control cell undergoing single-step vertical confinement, corresponding to Fig. 3 (a) and (b).** Confinement is applied at 00:50 (min:s) (10th time point; 5 s intervals), reducing the inter-gel spacing from 9.5 µm to 8 µm, as indicated by the schemes. The cell responds with the emergence of blebs beginning at 2:15 (min:s), with visible bleb expansion initiating at 4:15 (min:s) (white arrows). The accompanying stress data demonstrate a transient increase in lateral and normal stress following confinement, preceding bleb formation. Stress gradually stabilizes after the start of bleb expansion. Scale bar: 10 µm.

**Supplementary Video 12**

**Time-lapse of a DMSO-treated control cell exhibiting pre-existing blebs and undergoing bleb expansion.** Confinement is applied at 00:50 (min:s) (10th time point; 5 s intervals), reducing the inter-gel spacing from 7 µm to 6 µm, as illustrated by the schemes. The cell displays blebs at the onset of imaging and initiates bleb expansion shortly thereafter. The accompanying stress plot shows a gradual reduction in both lateral and normal stress over time, consistent with cellular adaptation to confinement. Scale bar: 10 µm.

**Supplementary Video 13**

**Time-lapse sequence of a Y-27632–treated cell undergoing single-step vertical confinement, corresponding to Fig. 4 (a) and (b).** Confinement is applied at 00:50 (min:s) (10th time point; 5 s intervals), reducing the inter-gel spacing from 11 µm to 6 µm, as shown in the schemes. Despite the strong confinement, no bleb formation occurs, consistent with ROCK inhibition. Lateral and normal stress levels remain minimal throughout the time course. A small, transient increase in stress is detected immediately following confinement, but no sustained stress development is observed. Scale bar: 10 µm.

**Supplementary Video 14**

**Supplementary video corresponding to Fig. 4 (e) and (f), illustrating the mechanical response of a blebbistatin-treated cell under single-step vertical confinement.** Due to the phototoxicity of blebbistatin under live-cell fluorescence, time-lapse imaging of the cell channel was not performed. Instead, representative top-view images from the cell before and after the time-lapse imaging are shown. The full time-lapse sequence is captured in the gel channel, where vertical confinement reduces the inter-gel spacing from 9 µm to 3 µm, as depicted schematically. Despite treatment, bleb formation is observed in the post-timelapse image, indicating that blebbistatin does not fully suppress blebbing under confinement. The accompanying plot shows lateral and normal stress over time for the same cell (Fig. 4e), demonstrating reduced stress levels in response to blebbistatin treatment compared to the control cells. Scale bar: 20 µm.

**Supplementary Video 15**

**Supplementary video corresponding to Fig. 6. Representative video of periodic confinement applied to a HeLa cell, with the inter-gel spacing reduced from 14.4 µm to 12 µm every 2.5 minutes during live imaging.** Both the bottom and top polyacrylamide (PAA) gels have a stiffness of 3 kPa. The red planes below and above the cell represent the surfaces of the bottom and top PAA gels, respectively. The thickness of the gels is 50–80 µm. Actin is labeled with LifeAct-GFP (green), and fluorescent beads embedded in the PAA gels are shown in red. Scale bar: 20 µm.

#### Supplementary Note 1

#### Technical design, dimensions and material specifications of the CFM device

Technical drawings in this Supplementary Note were prepared by the mechanical workshop in Münster.

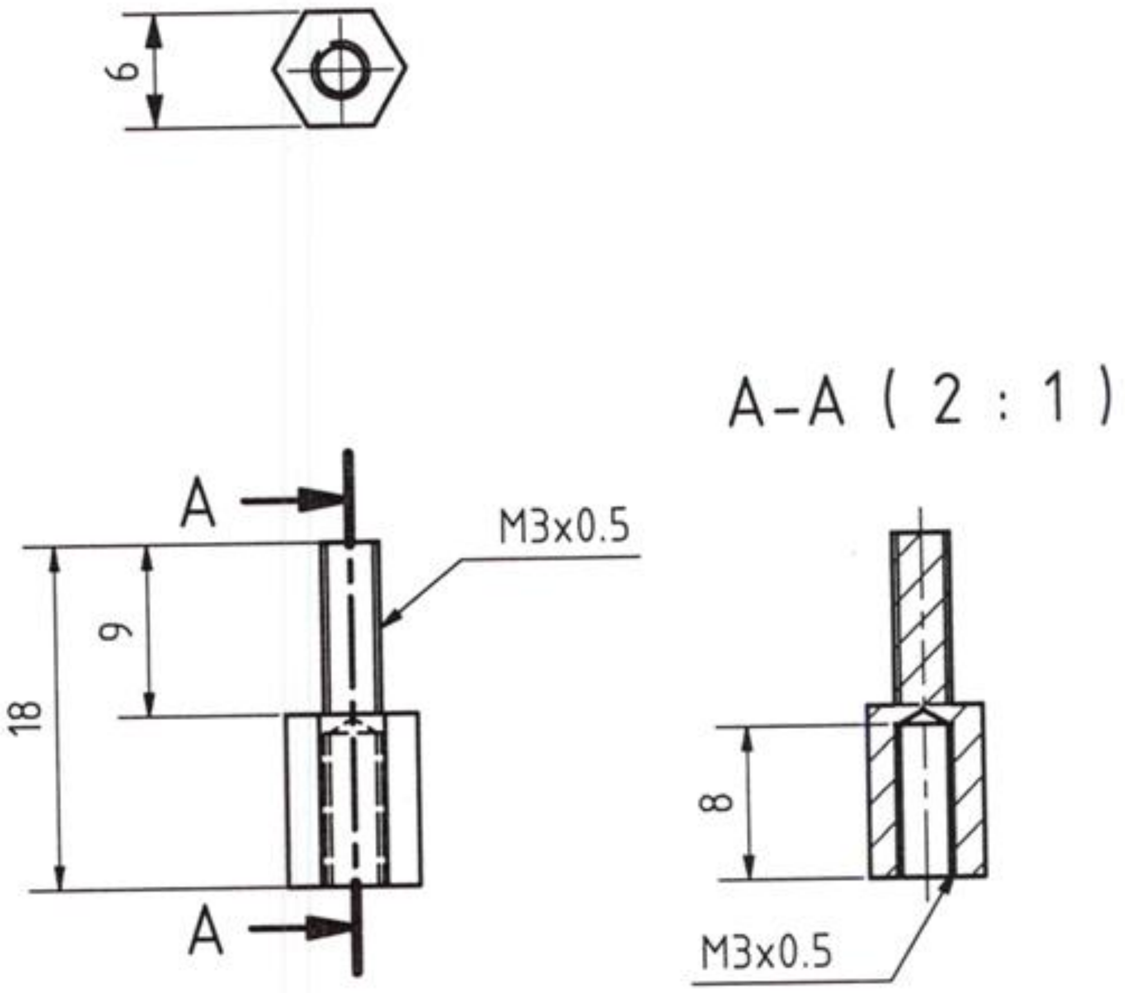
6
A
A
9
18
M3x0.5
A-A ( 2 : 1 )
8
M3x0.5

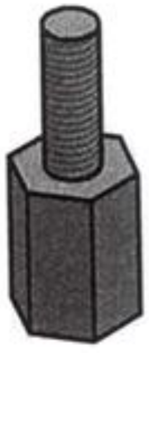

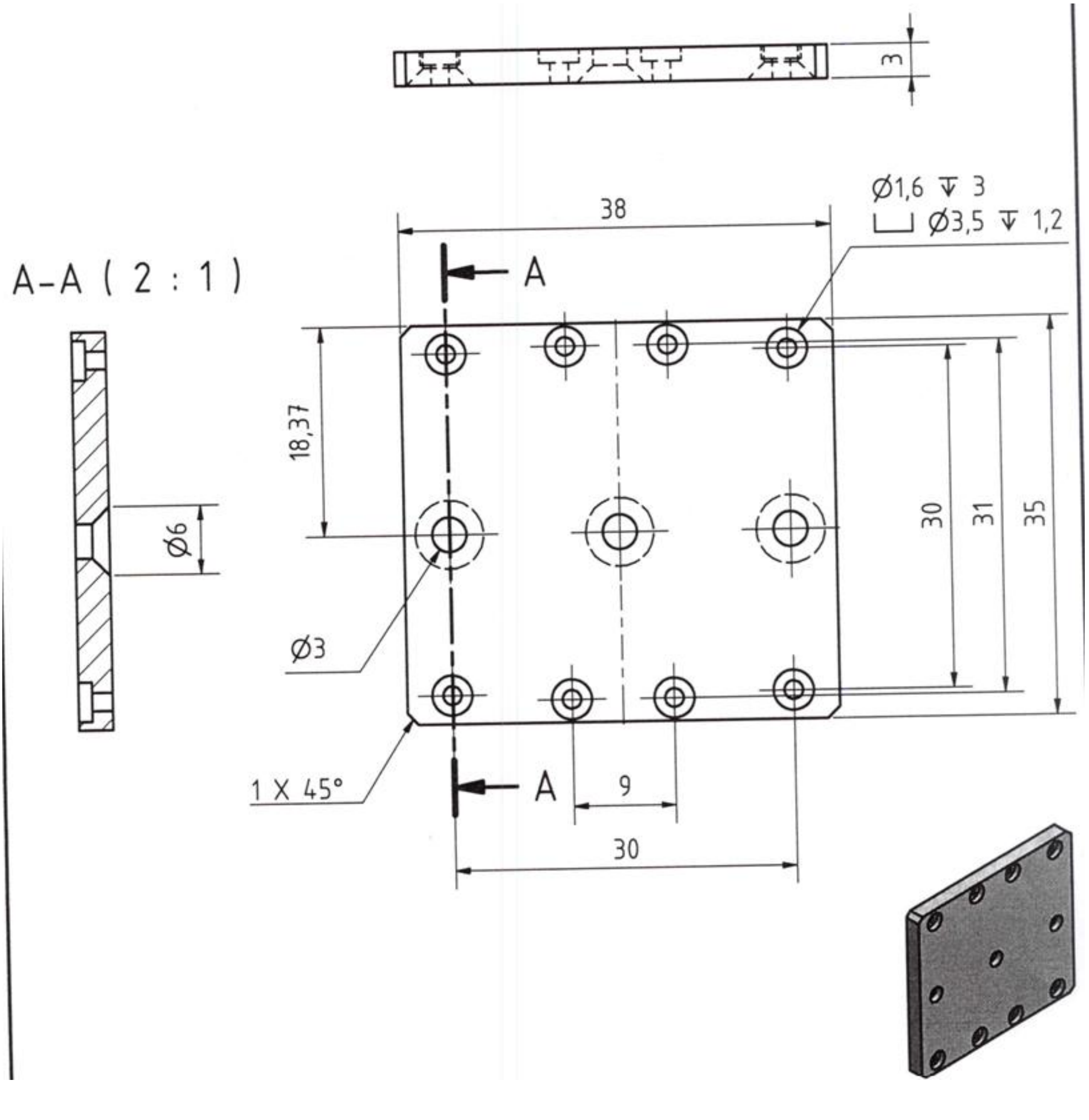
A-A ( 2 : 1 )
3
Ø1,6 ↧ 3
⌴ Ø3,5 ↧ 1,2
38
A
18,37
Ø6
Ø3
30
31
35
1 X 45°
A
9
30

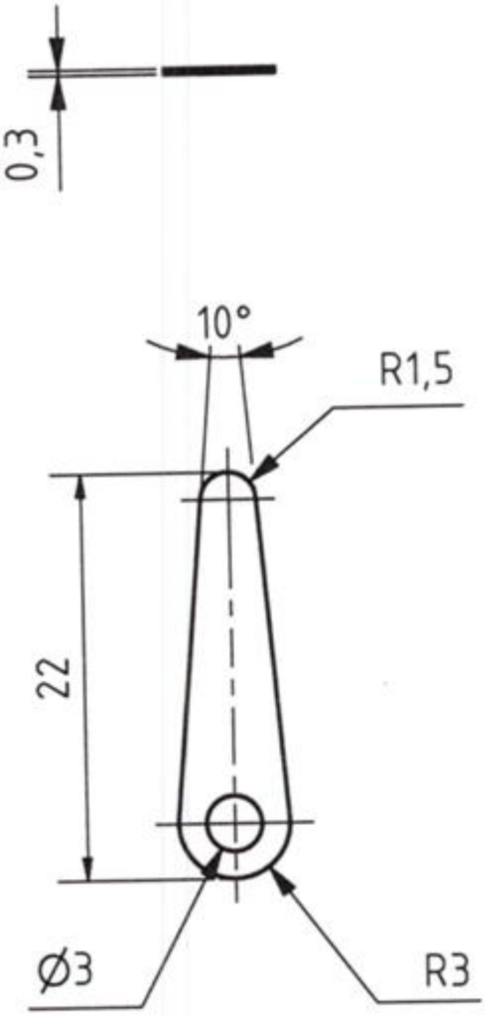
0,3
10°
R1,5
22
Ø3
R3

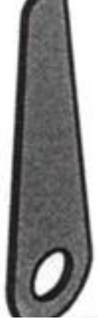

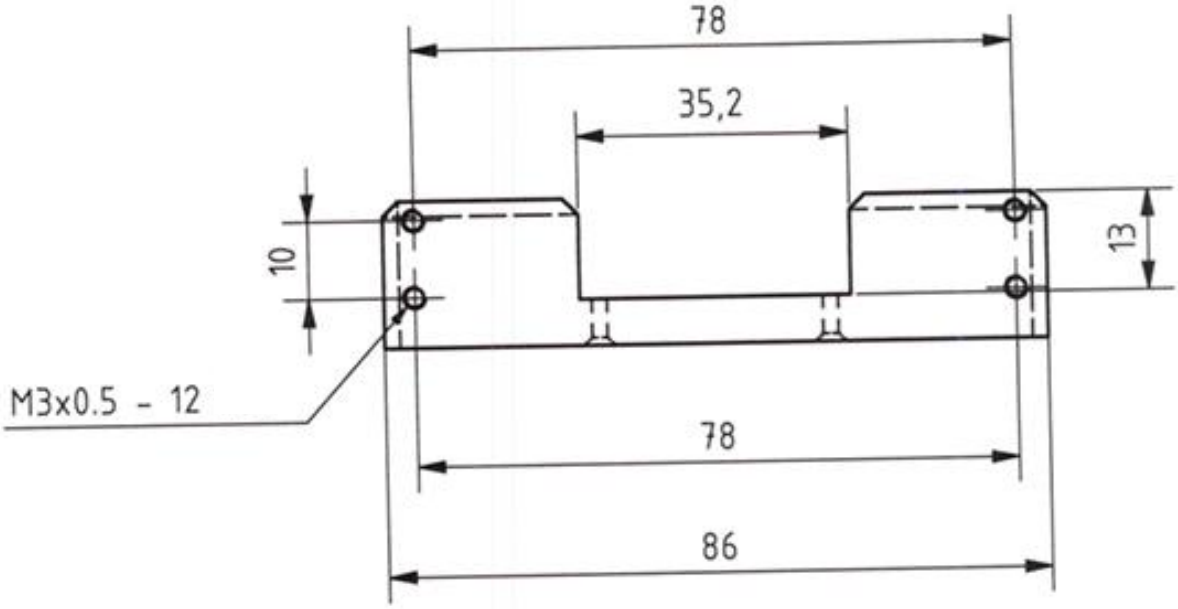


A-A ( 1 : 1 )

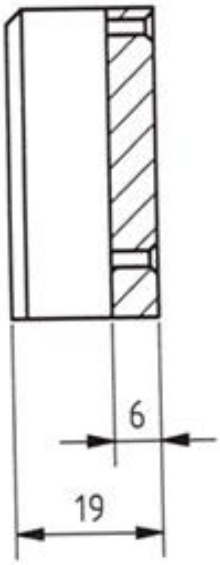


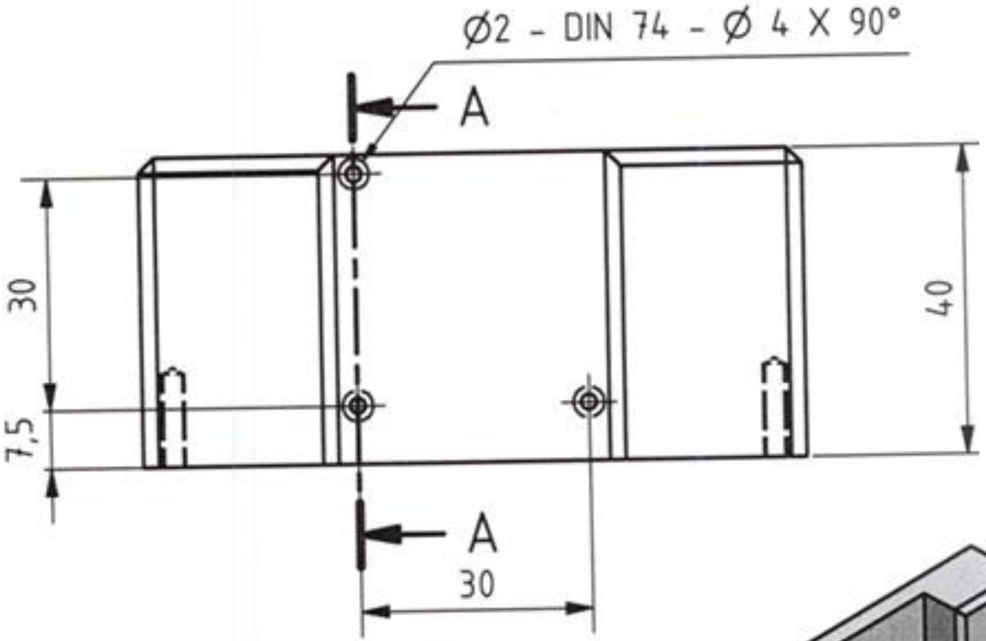


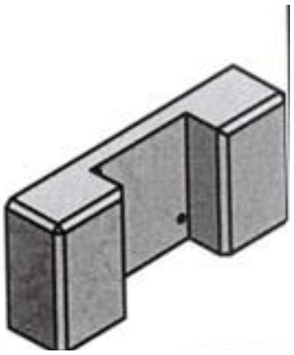

2 x45°

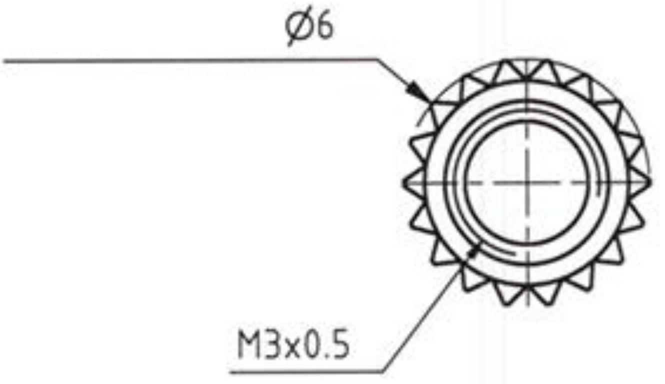
Ø6
M3x0.5

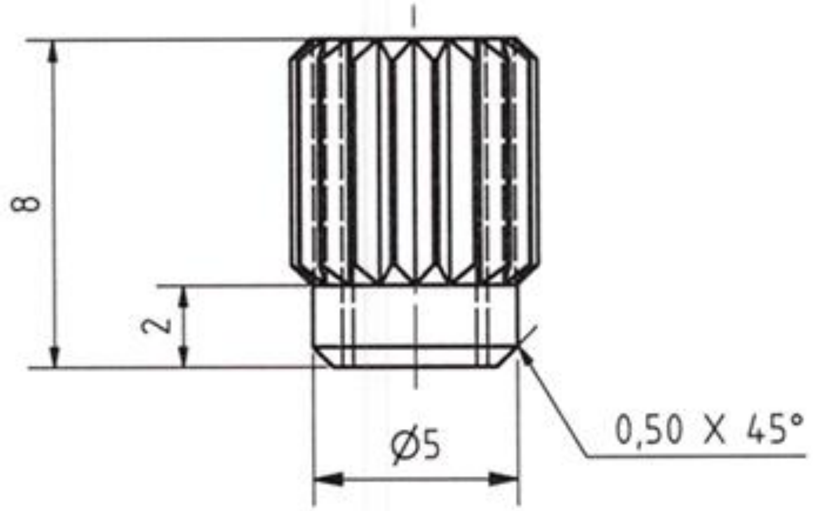
8
2
Ø5
0,50 X 45°

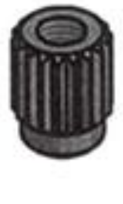

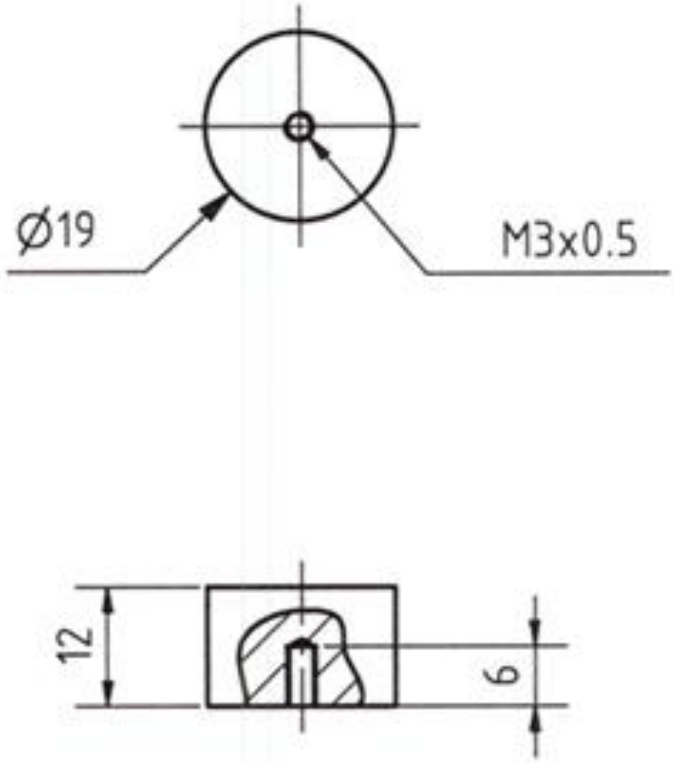
Ø19
M3x0.5
12
6

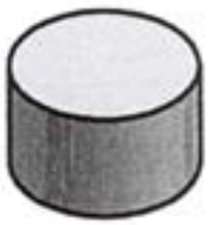

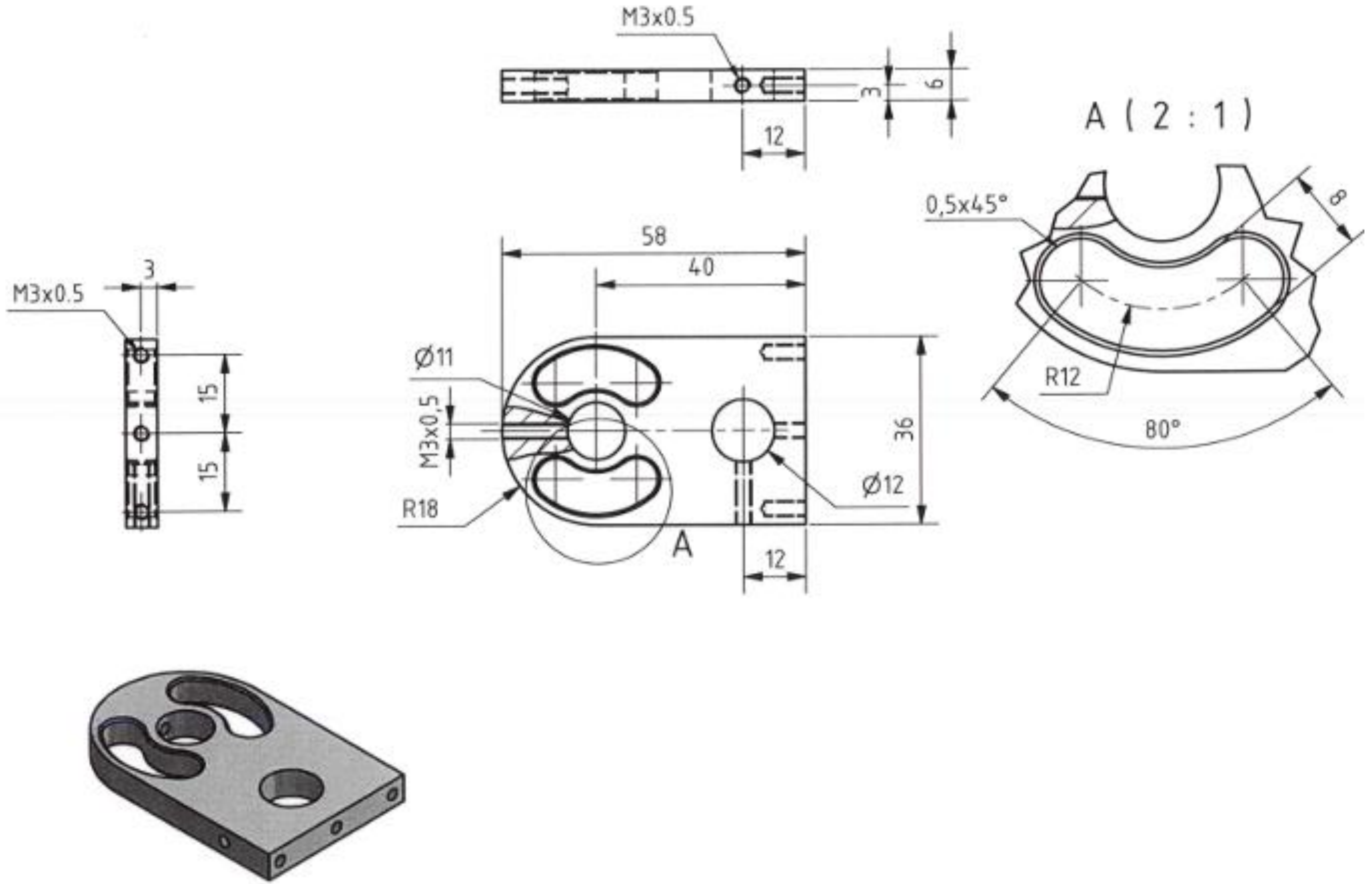
M3x0.5
3
6
12
A ( 2 : 1 )
0,5x45°
8
R12
80°
M3x0.5
3
15
15
58
40
Ø11
M3x0,5
R18
A
12
36
Ø12

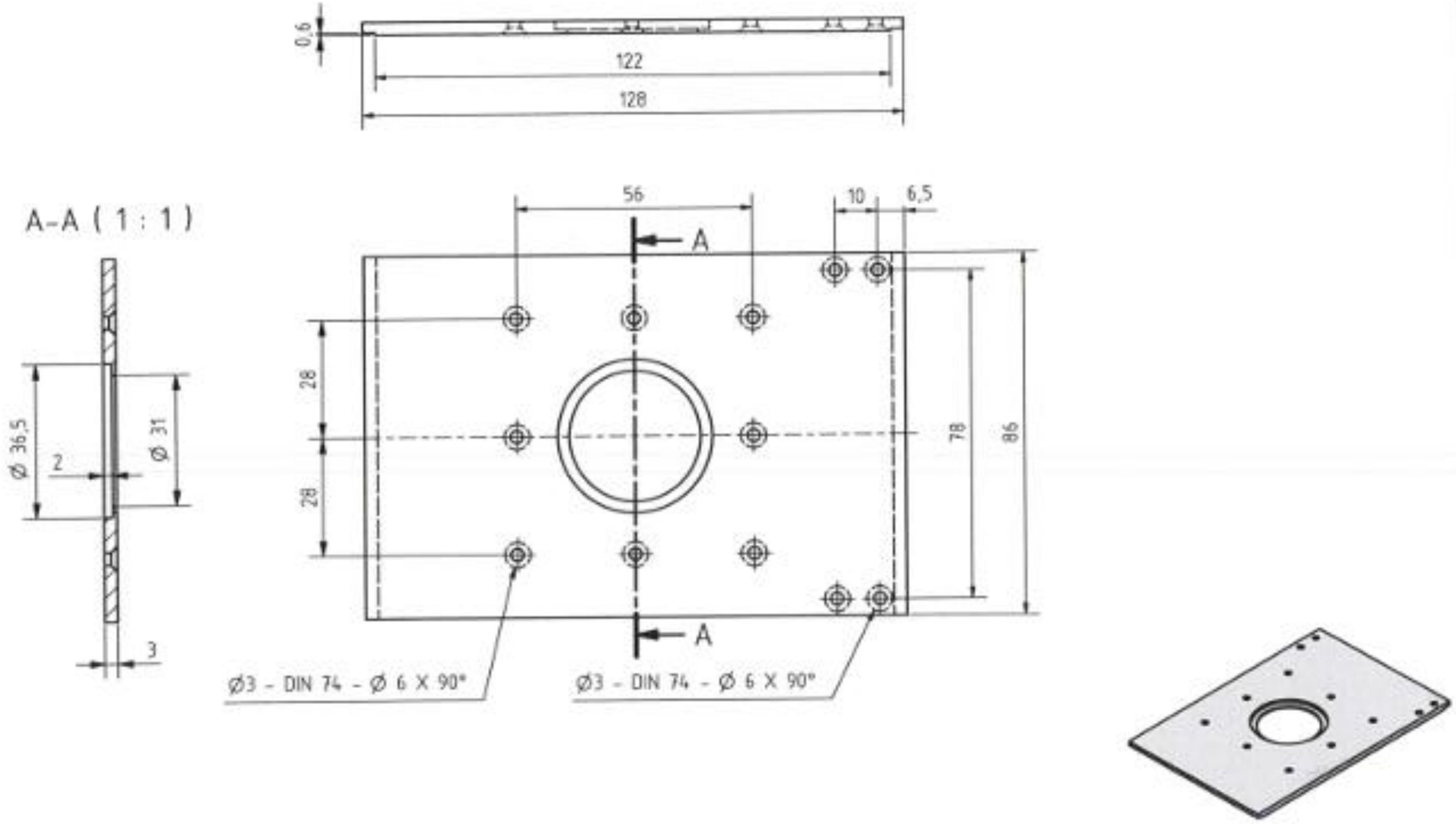
0,6
122
128
A-A ( 1 : 1 )
Ø 36,5
2
Ø 31
3
56
10
6,5
A
A
28
28
78
86
Ø3 - DIN 74 - Ø 6 X 90°
Ø3 - DIN 74 - Ø 6 X 90°

#### Supplementary Note 2. Effect of confinement level and microenvironment stiffness on neutrophil migration and its nucleus

Neutrophils are born in the bone marrow, and, to reach the site of infection, they have to move through very tight junctions and undergo extreme cellular and nuclear deformations, such as during transendothelial migration[20]. To study how these fast-migrating cells can undergo such extreme compression, a confinement instrument is required that can capture neutrophils' response immediately after confining them; otherwise, due to their very quick response to the external mechanical signals, it would not be possible to find out the mechanism behind their confined behavior. Using CFM, it is possible to confine these cells during live imaging and thereby capture their very early responses to external forces. To investigate how the microenvironmental stiffness affects the responses of neutrophils to the level of confinement, we varied both confinement and stiffness, and probed the cells' 2.5D traction stress in response to these changes (Supplementary Figure 1a-b and Supplementary Video 3). In order to simulate physiological *in vivo* conditions experienced by neutrophils, we exposed them to three different substrate stiffnesses, namely Young's moduli of 1.2, 9 and 21 kPa. In an initial step (Supplementary Figure 1a), we categorized our experimental design into two main groups: First, unconfined cells, seeded on a single PAA gel layer, and second, confined cells, seeded between two layers of PAA gels of the same stiffness. The confinement level was adjusted to a range of 5.4–7.8 $\mu$m (Supplementary Figure 1a). Our results demonstrated that independent of the microenvironmental stiffness applied, neutrophils in general exert higher lateral and normal stresses on their environment when confined. This suggests that neutrophils respond to the external confinement by an increase in pressure and traction forces. We also observed an increase in lateral and normal traction stress as a function of substrate stiffness both in unconfined and confined situations (Supplementary Figure 1a), which is consistent with the well-known effect of mechanotransduction, so far only studied in unconfined systems[21–23]. Lateral and normal stresses increased with increasing substrate stiffness but saturated in confinement already at 9 kPa substrate stiffness. Besides comparing confined and unconfined situations, we also tested the stress response upon increasing confinement levels at different substrate stiffnesses (Supplementary Figure 1b). To this end, we applied three different confinement ranges of 2.4–4.8 $\mu$m, 5.4–7.8 $\mu$m and 8.4–10.8 $\mu$m. We also used a greater distance (about 30 $\mu$m) between the gels as our control condition, which ensured a situation without contact between the neutrophils and the top gel, hence resembling a 2D substrate for the cells. We observed that increasing the confinement level led to an increase in both lateral and normal stress in a soft environment (1.2 kPa). In the medium and stiff environments (9 kPa and 21 kPa, respectively), the picture changed slightly as here both lateral and normal stress still increased for higher confinements, but after a critical confinement, below 5.4 $\mu$m, compressed neutrophils reversed their stress response, measured as a decrease in force at high confinements. Please note that the normal stress in the 9 kPa situation did not decrease significantly at high confinements, but still broke the trend of increased normal forces upon increased confinement as would be intuitively expected (Supplementary Figure 1b). In short, these results suggest that confined neutrophils adjust their pressure and lateral forces on the microenvironment as a strategy to cope with the external compression and difficulties on their way. The fact that such adjustment is not observed in HeLa cells points out the importance of studying such properties for a better understanding of cell-type-dependent mechanical response to confinement.

A notable feature of neutrophils under confinement is nuclear reorganization, which has been suggested to facilitate efficient migration even under strong confinement[24]. To further demonstrate the potential of CFM, we labeled the nucleus with a fluorescent live marker (SYTO™ Red Fluorescent Nucleic Acid Stain Sampler Kit, No. 60, Thermo Fisher) and tracked the nuclear shape changes in response to confinement. We observed noticeable nuclear deformation under high confinement (Supplementary Figure 1c; Supplementary Video 4), in contrast to the more rounded nuclear morphology in non-confined neutrophils (Supplementary Figure 1c; Supplementary Video 5). These deformations were especially evident when cells encountered another cell along their migration path (Supplementary Figure 1d; Supplementary Video 4). To test this further, we measured the surface area of the nucleus under different levels of confinement and compared it with the cell surface area. Surprisingly, we did not observe any significant change in the surface area of the neutrophil nucleus under different levels of confinement. In contrast, the cell surface area increased drastically for confinements below 5.7 $\mu$m (Supplementary Figure 1e). The multilobular nucleus is thought to facilitate neutrophil migration through narrow intercellular junctions due to its ability to undergo pronounced shape changes.

In short, CFM provides the possibility to correlate nuclear shape changes with force generation and migration characteristics, which will pave the way for new insights into active migration properties of immune and other cell types.

#### Supplementary Note 3. Neutrophil 3D durotaxis

Although 3D confinement is one of the major physical obstacles to cell migration in their natural environment, it remains difficult to study under controlled conditions. In addition, most migrating cells experience a range of microenvironmental stiffnesses while traversing different tissues. Neutrophils are a prime example: originating in the bone marrow, they migrate through small junctions and pores across tissues of diverse stiffness to reach sites of infection. Thus, along their journey they encounter tissue stiffnesses from $\sim$ 300 Pa (brain) to $\sim$ 10 GPa (bone)[25,26]. Neutrophils are also responsive to substrate stiffness, adjusting adhesion and spreading[27]. Potential stiffness gradients can therefore bias confined migration (durotaxis), a phenomenon well documented in 2D culture. Confinement Force Microscopy (CFM) enables us to study 3D durotaxis of these rapidly moving cells under confinement, avoiding the 2D in-gel gradients typical of classical assays. By preparing two PAA gels of distinct stiffnesses and positioning them above and below the cells, we can precisely trigger the moment when neutrophils encounter a well-defined stiffness contrast, allowing us to capture the immediate response to confinement and steep stiffness gradients. Here, we confine neutrophils between two PAA layers of different stiffnesses (bottom: 1.2 kPa; top: 114 kPa) while maintaining a constant gap for the desired imaging period until a complete durotactic switch from the soft to the stiff substrate is observed (Supplementary Figure 2a; Supplementary Video 6). By registering the traction stresses on the soft substrate with the shifting process, we observe a surprising transient stress peak on the soft substrate during neutrophil durotaxis (Supplementary Figure 2b). This peak occurs when neutrophils adopt a cylindrical configuration between the substrates (Supplementary Figure 2a; at 5:33 min:s, the stress peak occurs), a moment when the attachment area is approximately equal on the soft and stiff sides. Estimating forces from stress and the cell–gel contact area indicates that the total force on the soft side remains nearly unchanged from the initial state (cell fully spread on soft) to the cylindrical state. Shortly after this equal-area moment, forces on the soft gel drop sharply as cells spread on the stiff substrate and detach from the soft (Supplementary Figure 2c). This behavior may arise from focal adhesion and/or cytoskeletal reorganization during durotaxis.

We also tested the inverted configuration (soft top $\rightarrow$ stiff bottom; top 1.2 kPa, bottom 114 kPa; Supplementary Video 7) and observed robust transitions. However, we could not quantify tractions on the stiff bottom: neutrophils are weakly adhesive and transition rapidly (sometimes $< 3$ min), yielding no resolvable bead displacements in 114 kPa gels, so the TFM inversion returned no stress signal. Lowering the “stiff” substrate to $\sim$50 kPa increased sensitivity but abolished durotaxis within our 30 min observation window.

#### Supplementary Note 4. Measuring force generation during development

Embryogenesis involves a complex interplay of cellular and molecular processes that shape the developing organism. While embryogenesis is governed by tightly regulated genetic programs, it is equally dependent on mechanical forces, which drive tissue morphogenesis to ensure robust development. Perturbation of these forces can disrupt the entire process, leading to numerous developmental defects. However, our understanding of their roles remains limited, largely due to technical challenges in measuring forces *in vivo*. To demonstrate the power of confinement force microscopy (CFM) in classical developmental model systems, we confined developing *Drosophila melanogaster* embryos and monitored force generation during cellularization—the first major tissue formation step, which depends on membrane-coupled actin remodeling and myosin-driven contractility[28]. Using Squash-GFP–expressing embryos (Drosophila myosin II homolog), we imaged embryos just prior to cellularization (Supplementary Figure 3a; Supplementary Video 8). Despite confinement, embryos progressed through cellularization without detectable defects. Notably, deformation of the underlying PAA gel revealed a small but measurable increase in normalized displacement as the actomyosin ring detached from the membrane (Supplementary Figure 3b), consistent with increased pushing forces against the surrounding matrix during cellularization.

#### Supplementary Note 5. Measuring stresses imposed on the microenvironment during cell death

Apoptosis, or programmed cell death, is vital for the tissue morphogenesis and the elimination of unnecessary cells during embryonic development, tissue homeostasis, and certain pathological conditions[29]. The mechanical forces produced during apoptosis are important not only for extruding dying cells from tissues to maintain tissue integrity but also for altering the morphology of neighboring cells to fill the space originally occupied by the dying cell[30]. These forces could also play a role in other biological processes, such as the tissue-fusion process (dorsal closure) during *Drosophila* embryogenesis[31] or hair follicle regression[32].

To test the capabilities of CFM for studying the general process of cell death, we imaged C2C12 cells (precursor muscle cells) that were confined between PAA gels with stiffnesses of 1.2 and 21 kPa for the bottom and top gels, respectively. The C2C12 cells were stained with CellMask™ Deep Red plasma membrane stain, which is known to be toxic to cells under high-power laser illumination[16]. Cells were kept under confinement for about 12 hours and exposed to the high-power imaging laser every 30 min. During early cell death, cells reduced their projected area while simultaneously increasing sphericity, thereby indenting the gels; this preceded plasma-membrane rupture and subsequent disintegration (Supplementary Figure 4a-b; Supplementary Video 9). Because the bottom gel had ∼20-fold lower stiffness than the top gel, cells were able to deform it readily to make space for their increased height. The bottom-gel lateral and normal stress traces consistently peaked at the time of maximal deformation and then decayed to near zero (Supplementary Figure 4c), consistent with loss of active force generation upon death. Supplementary Figure 4c-d show the average lateral and normal stresses across cells, normalized to each cell's minimum bottom-gel stress. On average, the lateral stress increased by ∼2-fold and the normal stress by ∼5-fold around the death event.

#### Supplementary Note 6. Measuring the viscoelastic properties of cancer cell spheroids

Viscoelasticity is a fundamental property of cells and tissues and varies across cell lines and tissue types[33–37]. The viscoelasticity of the resident tissue could be a determining factor in cell fate[38,39]. Likewise, tumor viscoelasticity could affect the phenotype and proliferation of malignant cells[40]. Therefore, viscoelasticity is an important feature for distinguishing healthy from malignant tissue. Accurate measurements of viscoelasticity could thus be instrumental in the early diagnosis of cancer, with the potential to save many lives. In this regard, we used CFM to measure the elasticity and viscosity of cancer spheroids formed by different cancer cell lines. By confining HeLa and CT26 spheroids while recording the spheroids' strain and stress relaxation over time (Supplementary Figure 5a-c; Supplementary Video 10), we obtained stress–strain relations. These can be explained by a mechanical Maxwell model (Supplementary Figure 5c), which fits the relaxation data well and allows calculation of elastic and viscous properties. Our results demonstrate that although the stiffnesses of CT26 spheroids and HeLa spheroids are not significantly different, CT26 spheroids are considerably more viscous than HeLa spheroids (Supplementary Figure 5d–e). Supplementary Video 10 illustrates this very clearly. It can be seen that HeLa cells move out of the spheroid and the spheroid behaves more fluid-like than the CT26 spheroid, where the superficial cells remain strongly adherent to the spheroid and surface tension appears higher. Our results are highly consistent with micropipette aspiration measurements from previous studies[41]. These results demonstrate that CFM might be a suitable alternative to complicated systems such as micropipette aspiration.

#### Supplementary Note 7. Statistical association between mechanical and morphological responses under confinement

To relate mechanics to morphology under vertical confinement, we computed Pearson correlations across key single-cell variables. Mechanical variables were obtained from the gel-channel time series, whereas blebbing before and after imaging was scored from cell-channel images acquired immediately before and immediately after the time-lapse. Only cells for which all variables could be scored unambiguously were included in this analysis ($n = 74$ cells; Supplementary Figure 6). Variables were encoded either as binary/categorical variables or as continuous variables. Binary variables were coded as yes = 1 and no = 0. Lateral- and normal-stress trends were coded as increase = 1, no clear change = 0, and decrease = -1. Pearson's correlation coefficient ($r$) quantifies the strength and direction of a linear association between two variables ($r = 1$, perfect correlation; $r = -1$, perfect anti-correlation; $r = 0$, no correlation).
Our results reveal the following relationships among key variables:
**Higher confinement strain increases the likelihood of observing an immediate post-step stress jump in both lateral and normal components.** Higher imposed strain (i.e., more compression) was associated with an increased likelihood that an immediate stress jump was detectable after the confinement step in both the lateral and normal directions ($r = 0.38$ and $r = 0.40$, respectively), consistent with a passive, geometry-driven load transfer at the confinement step.
**Prior confinement increases the probability of blebbing and is associated with a larger immediate normal-stress jump.** Cells that had already been confined at the start of imaging were more likely to bleb ($r = 0.41$) and to exhibit a detectable normal-stress jump ($r = 0.41$), suggesting a "preconditioning" effect whereby prior confinement increases cortical prestress and intracellular pressure; similar preconditioning effects have been described in other systems where prior mechanical exposure and prestress alters cytoskeletal or nuclear behavior[42–47].
**Stronger coupling between lateral and normal stress dynamics is associated with a decreasing normal-stress trend over time.** We quantified, for each cell, how tightly lateral and normal stresses rise and fall together over time ("corr.(lat&norm)", Pearson's $r$; Supplementary Figure 6). In general, lateral and normal stresses tended to evolve together ($r = 0.40$), indicating coordinated adaptation across orthogonal axes, although this coupling is heterogeneous at the single-cell level (only $\sim$50% show matched trends; Figure 2f).
More interestingly, cells showing stronger coupling between these two channels exhibited a decreasing normal-stress trend over time ($r = -0.56$), consistent with active adaptation that relieves vertical pressure via coordinated in-plane contractility and cortical remodeling[48,49]. Conversely, in cells where normal stress increases over time, this buildup is not consistently mirrored by lateral stress changes—implying a decoupling between stress components. Such asymmetry could reflect ineffective or delayed mechanical adaptation, or localized vertical force accumulation without sufficient redistribution across the cortex. These findings highlight the value of capturing both lateral and normal stresses to uncover force coordination strategies used by cells during confinement-induced adaptation.
**Pre-step blebbing is associated with a decreasing normal-stress trend over time.** Prior blebbing predicted subsequent stress relief ($r = -0.40$), supporting the view that blebbing functions as a pressure-release mechanism under confinement.

Together, these correlations reveal distinct mechanical and morphological signatures that define how cells respond to confinement. By simultaneously measuring orthogonal stress components and mapping them against confinement history and morphological changes, CFM enables a comprehensive understanding of mechanical adaptation at the single-cell level.
